# Chemotherapy-induced multicellularity drives drug-tolerant persistence state in tumor cells

**DOI:** 10.64898/2026.07.09.737606

**Authors:** Jing Li, Zefan Zhu, Entong Zheng, Ji Xiong, Aiping Liu, Tingting Hu, Zhan Ma, Chunfang Liu

**Affiliations:** Department of Laboratory Medicine, Huashan Hospital, Shanghai Medical College, Fudan University, Shanghai, 200040, China; Department of Pathology, Huashan Hospital, Shanghai Medical College, Fudan University, Shanghai 200040, China; Department of Laboratory Medicine, Shanghai Children’s Hospital, Shanghai Jiao Tong University, Shanghai 200040, China

**Keywords:** Chemotherapy-induced multicellularity, drug-tolerant persistence (DTP) state, phenotypic plasticity, tumor evolution, adaptive life-cycle alternation

## Abstract

Multicellularity is a well-documented microbial response to stress, however its role as an adaptive survival strategy in cancer remains unresolved. Here we reveal that drug stress, such as paclitaxel treatment, enable rapidly (within 24-48 hours) and efficiently (∼20-40%) convert single mouse breast 4T1 cancer cells into clonal multicellular spheroids, ultimately generating multicellular masses. Notably, multicellularity is reversible: upon stress removal, most of them restore a unicellular lifestyle that quickly becomes dominant. This transient multicellular state shields cells from hostile niches, functions as a drug-tolerant persistence (DTP) reservoir, and fuels post-therapy relapse, revealing multicellularity as a facultative evolutionary pivot for fitness gain. Importantly, blocking primordial germ cell (PGC) specification suppresses the multicellularity transition. Our findings reveal that certain cancer cells enable adopt unicellular–multicellular life cycle through phenotypic plasticity, dynamically adapting to microenvironmental shifts to maximize fitness. This discovery reframes cancer evolution and the drug-tolerant persistence (DTP) state, highlighting multicellularity as an adaptive, stress-inducible survival strategy against therapy.

**Highlight:**

1. Drug induced evolution from unicell to clonal multicellularity.
2. Providing a novel DTP state.
3. Revealing environment-dependent bistable plasticity: multicellular persistence (stress) versus unicellular proliferation (favorable).
4. Blocking primordial germ cell (PGC) specification suppresses the multicellularity.

## Instruction

Recent oncology data reveal that a principal driver of cancer relapse is not the acquisition of resistance-conferring mutations, but rather the reversible, non-genetic “drug-tolerant persistence” (DTP) state that provides only transient protection^1,2^. Strikingly, microorganisms can be rapidly switched by microenvironmental stress into multicellular lifestyle, whether aggregative biofilms or clonal cysts^3–6^, as a core adaptive strategy that transiently buffers them against lethal antibiotic exposure. We propose that cancer cells may adopt an analogous program to enter a DTP state. Here, we report that chemotherapeutic stress alone—typified by paclitaxel (PTX)—is sufficient to drive a reversible unicellular-to-clonal-multicellular transition in a fraction of malignant cancer cells, often characterized by the co-emergence of oocyte-like and blastomere-like phenotypes, thereby driving cells into a robust DTP state. Our findings redefine the nature of the DTP state and frame cancer adaptive evolution as the re-activation of an ancient developmental program.

### Drugs induced unicellular-to-multicellular transition in cancer cells

To investigate the induction of unicellular-to-multicellular transition under therapeutic stress, we established a model in which the 4T1 cultures was exposed to sustained drug stress, including PTX, vincristine (VCR), doxorubicin (Dox), cisplatin (DDP) and 5-Fluorouracil (5-FU). Under standard culture conditions, 4T1 cells proliferated robustly and remained predominantly unicellular (Figs. 1a and 1e). And ∼99 % of round cells were <20 µm in diameter (12–15 µm peak) whereas large round cells (>20 µm in diameter) were rare, accounting for only ∼0.1% (Figs. 1a and 1b, Tables S1 and S2). Interestingly, after treatment with 10 µM PTX or VCR, the population of large round cells (>20 µm in diameter) increased markedly (Figs. 1a-c, Tables S1 and S2). Notably, the large round cells—whether mononuclear or multinuclear—displayed pronounced drug tolerance: a fraction not only survived but continued to expand during prolonged culture, ultimately generating multicellular state, even large multicellular mass (Figs. 1a, 1c-f and S1a). By contrast, standard cultures never produced such giant multicellular masses, indicating that drug stress alone can trigger multicellularity in certain cancer cells (Figs. 1a-f and S1a, Tables S1 and S2, S-Movie 1-4).

**Fig. 1.**
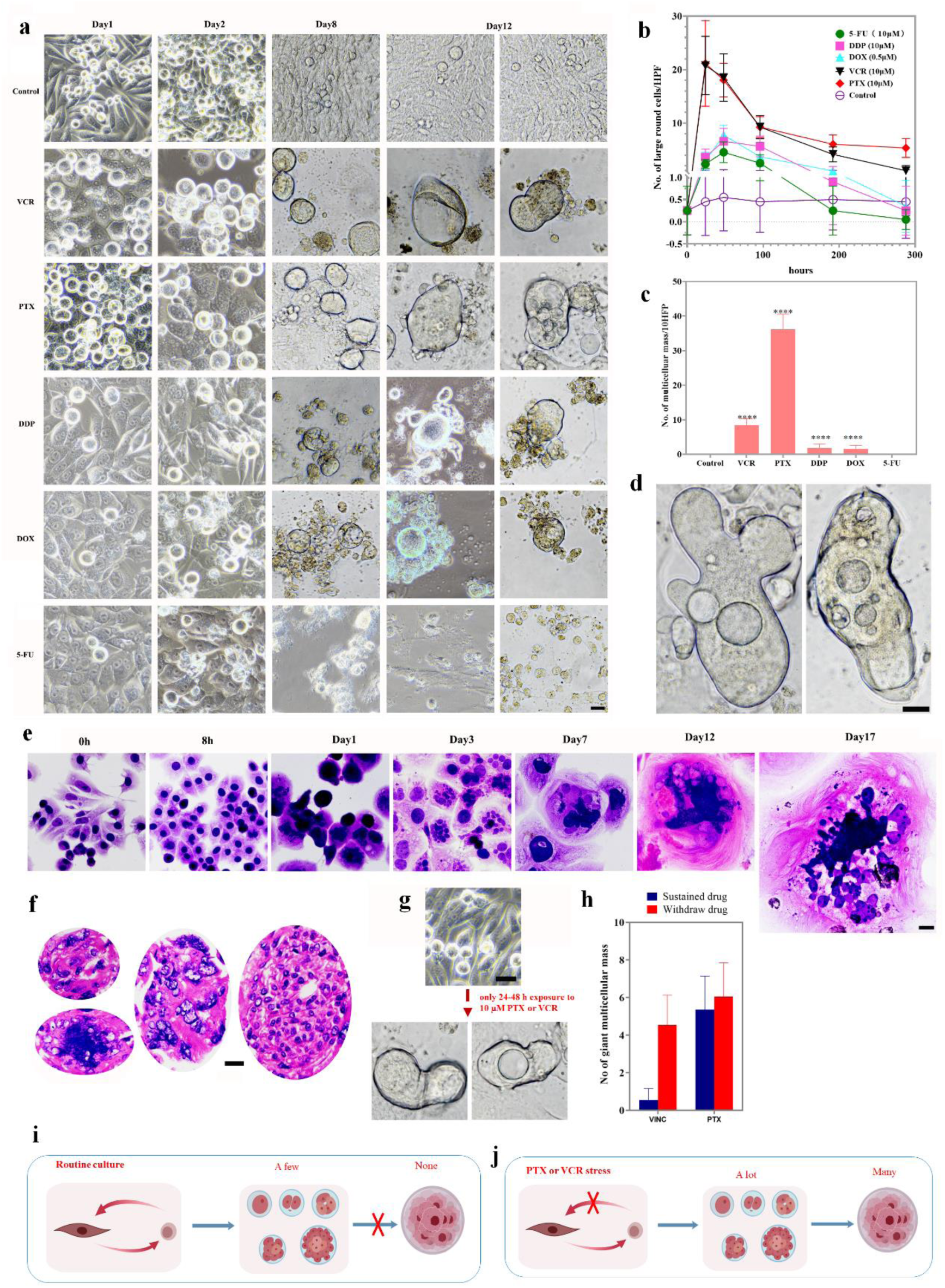
Drug-induced multicellularity in 4T1 cultures. (a) Bright-field images of cell morphology over time under control conditions versus exposure to 1 µM doxorubicin or 10 µM PTX. (b, c) Quantification of large round-cell formation capacity (n = 20) and multicellular mass counts after 12 days of treatment. (d–f) Representative images of multicellular aggregates after 18-day continuous PTX exposure, with corresponding H&E-stained sections shown in (e, f). (g, h) Pulse-chase experiments: 24–48 h exposure to 10 µM PTX or VCR followed by drug-free culture triggers giant multicellular mass formation; comparison of 48 h drug pulse versus continuous 12-day treatment. (i, j) Schematic models of 4T1 cell-fate trajectories under standard conditions (i) versus doxorubicin/PTX stress (j). Scale bar = 25 μm. Statistical significance indicated as *P < 0.05, **P < 0.01, ***P < 0.001, ****P < 0.0001.

Our data revealed that drugs (PTX and VCR) drove multicellularity far more potently than other agents (Dox, DDP and 5-FU) (Figs. 1a-c, Tables S1 and S2). PTX and VCR, rapidly and robustly induced the increase of cell size, whereas Dox, DDP and 5-FU did so slowly and at markedly lower efficiency (Figs. 1a-c, Tables S1 and S2). Compared with the control, the number of large round cells (>20 µm) rose dramatically after 24 h of drug exposure, increasing almost 100-fold in both the PTX- and VCR-treated groups (Figs. 1a-c, Tables S1 and S2). PTX and VCR produced significantly more giant multicellular masses than the other agents (Figs.1a and 1b, Tables S1 and S2). Although the two drugs were equally efficient in inducing large round cells, PTX was markedly more potent in driving the formation of giant multicellular masses (Figs. 1b and 1c, Tables S1 and S2). Notably, after only 24-48 h exposure to 10 µM PTX, followed by complete drug removal, 4T1 cells still executed the unicellular-to-multicellular transition and generated giant multicellular masses (Fig. 1g), revealing that a brief PTX pulse is sufficient to irreversibly trigger multicellularity. Additionally, brief drug exposure followed by washout markedly enhanced the survival of induced large rounded cells, consequently increasing the number of giant multicellular masses in the VCR-treated group (Fig. 1h, Table S3). Together, our findings reveal that clinically relevant cytotoxic stress drives a unicellular-to-multicellular transition in specific cancer cells (Figs. 1i and 1j), uncovering a shared adaptive evolution triggered by distinct chemotherapeutic agents.

### Essential role of multicellularity in survival

After 12 days of continuous drug exposure, the majority of cells died, while a small population remained viable (Fig. 2a). Of note, the majority of viable cells had transitioned into a multicellular state (Fig. 2b). The drug’s heightened capacity to generate multicellular-state cells was associated with an enrichment of survival cells (Figs. 1b and 2a). Because PTX most efficiently triggered the unicellular-to-multicellular transition among the tested drugs (Figs. 1a and 1b; Table S4 and S5), we selected it as a model to further define the contribution of multicellular state to survival. After 21 days of continuous 10 µM PTX exposure, the survival cells were overwhelmingly multicellular: ∼15 % multicellular spheroids and ∼80 % multicellular masses, whereas mononuclear small and large round cells accounted for only ∼0.5 % and ∼4.5 % of the survivor pool, respectively (Fig. 2c). H&E staining revealed that the majority of these 21-day PTX-treated masses exhibited a multicellular histological morphology (Figs. 2d and S1b). The findings indicated that multicellularity is the predominant DTP state.

**Fig. 2.**
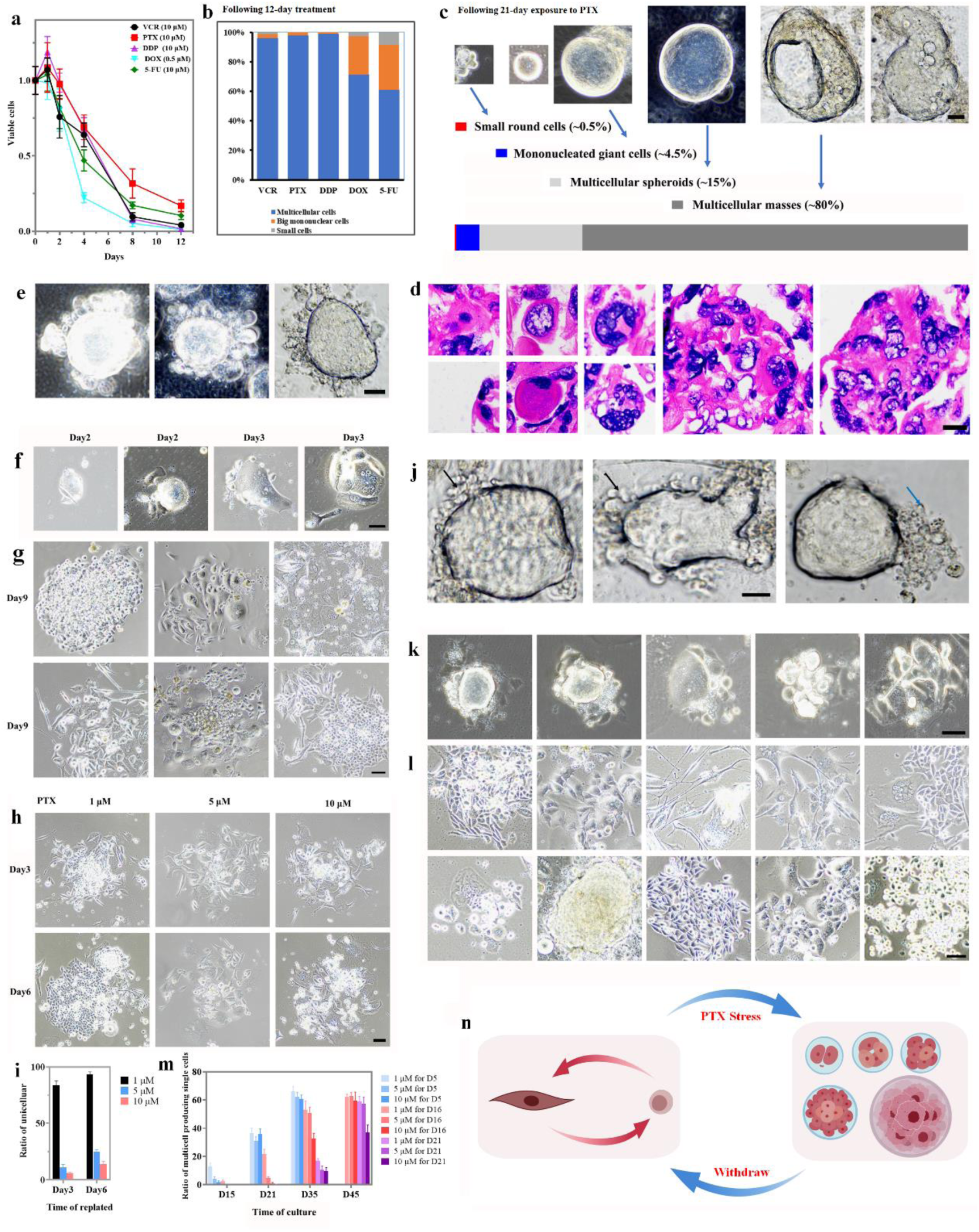
Bidirectional unicellular–multicellular transitions in 4T1 cultures. (a–c) Cell viability time-course (n=8) and survival rates following 12- and 21-day treatment with indicated drugs. (d) H&E-stained sections of 21-day PTX-treated cultures. (e, j) Daughter cells (black arrows: viable; blue arrows: PTX-killed/collapsed) surrounding multicellular spheroids in PTX-treated cultures. (f–i) Spheroids from 4–6 day PTX-treated cultures were suspended and replated in drug-free medium to generate daughter cells at indicated timepoints. (k, l) Daughter cells from PTX-treated (5 µM, 6 days) cultures after 4- or 15-day drug withdrawal. (m) Quantification of daughter cell generation capacity from drug-treated multicellular cells following various withdrawal periods. (n) Schematic model of microenvironment-driven bidirectional transitions. Scale bar = 25 μm. Statistical significance: *P < 0.05, **P < 0.01, ***P < 0.001, ****P < 0.0001.

### Environment-dependent unicellular–multicellular bistable plasticity

We then investigated whether the unicellular and multicellular states in 4T1 cultures could mutually interconvert in response to environmental changes. Under sustained PTX pressure, early multicellular spheroids could initially seed surrounding progeny (Fig. 2e), yet the bulk of these colonies collapsed within 3–6 days of PTX exposure as daughter cells underwent cell death to drug stress (Fig. 2e). It is noteworthy that a subset of multicellular spheroids remained loosely adherent. Upon re-plating in drug-free medium, most of these loosely adherent spheroids rapidly re-attached and re-established proliferative colonies (Figs. 2f-h). The probability of establishing such daughter-cell outgrowths scaled inversely with both PTX dose and exposure time (Figs. 2h and 2i, Table S6). Upon drug withdrawal, >80 % of suspension-phase multicellular spheroids (pre-treated with 1 µM PTX for 6 day) re-attached and spawned daughter cells within 3-7 days ((Figs. 2h and 2i, Table S6). The capacity of loosely adherent spheroids/masses to revert to a unicellular state became progressively impaired with prolonged drug treatment beyond 8 days. Notably, a subpopulation of larger multicellular masses also shed progeny in vitro, yet the vast majority of these offspring perished under continued PTX pressure (Fig. 2j). Upon drug withdrawal, the proportion of persisting cells re-entering the cell cycle was >50% following 16 days of continuous 5 µM PTX exposure, compared to >30% following 21 days of continuous 10 µM PTX exposure (Figs. 2k-m, Table S7). Thus, the multicellular state is reversible, establishing a recurrent life cycle upon PTX stress and withdrawal: unicellular → multicellular → unicellular (Fig. 2n). We define this phenomenon as unicellular–multicellular bistable plasticity, reflecting the discrete, switch-like transitions between these two stable states. Collectively, our findings indicate that environmental stimuli can trigger this bistable life cycle program in specific cancer cell populations (Fig. 2n), establishing stress-inducible adaptive evolution as a fundamental mechanism underlying cancer cell plasticity.

### Lead to tumor relapse

Regardless of whether they originated from short-term drug-exposed suspension cultures or long-term drug-exposed adherent cultures, the daughter cells displayed heightened morphological plasticity (Figs. 2g and 2i) and retained robust tumorigenicity, forming embryonic, highly plastic tumors—including teratomas (Figs. 3a-3d, S2a-c). *In vivo*, 4T1 cultures pre-treated for only 2 days formed tumors rapidly, although subsequent growth was markedly slower (Figs. 3e and 3f, Table S8). Conversely, cultures pre-treated for 14 days remained tumorigenic, but tumor initiation were significantly delayed (Figs. 3e and 3f), similar to the clinical lag that often precedes relapse after chemotherapy^1,2^. The tumorigenic potential of 4T1 cultures declined in proportion to PTX exposure time (Figs. 3e and 3f, Table S9). H&E sections revealed that drug-treated explants retained the histologic morphology of control tumors (Fig. 3g), yet displayed amplified plastic features (Fig. S2d), consistent with the heightened plasticity frequently seen in clinical relapse specimens^7^. Collectively, our findings reveal that the multicellular state enables reversion to a unicellular lifestyle and thereby drives post-therapy relapse.

**Fig. 3.**
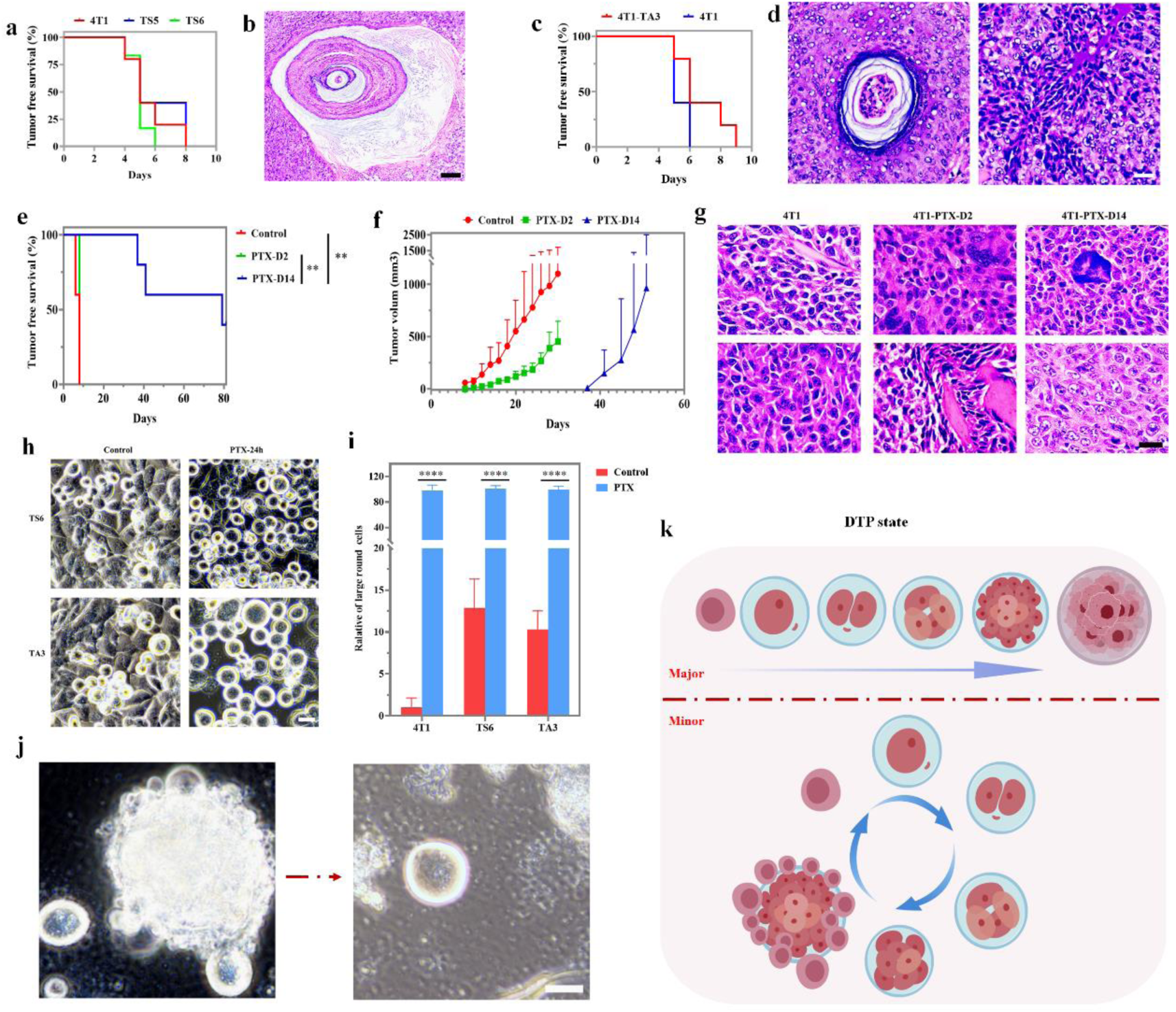
DTP states driving tumor relapse in 4T1 cultures. (a–d) Tumorigenicity and H&E histology of daughter cells derived from suspended spheroids (a, b) or drug-surviving cultures (c, d) (n=5). (e–g) Tumor initiation, growth kinetics, and pathological features in control versus PTX-pretreated 4T1 cells (n=5). (h, i) Phenotypic plasticity of daughter cells upon PTX re-challenge after drug withdrawal. (j) Rare persister cells from drug-induced spheroids surviving and proliferating under sustained PTX pressure. (k) Schematic model of novel DTP states facilitating tumor relapse. Scale bar = 25 μm (h, j), 50 μm (d, g), 100 μm (b). Statistical significance: *P < 0.05, **P < 0.01, ***P < 0.001, ****P < 0.0001.

### Novel DTP states

Recent studies have established non-genetic plasticity as a cornerstone of the DTP state and the subsequent evolution of stable resistance. DTPs are classically defined by four hallmarks^1,2^: (i) phenotypic adaptation independent of genetic alteration; (ii) deep quiescence or low cycling; (iii) intrinsic or drug-induced capacity to survive lethal drug concentrations; and (iv) full reversibility upon drug withdrawal. Consistent with DTP state, daughter cells recovered after PTX withdrawal remained drug-sensitive and rapidly re-entered multicellular state upon re-exposure (Figs. 3h and 3i, Table S10), revealing that these states confer only transient survival advantage without early acquisition of resistance mutations. Our data reveal that the induced large round cells and multicellular-masses meet the defining criteria of DTPs. Consequently, we identify the large round mononuclear cells, multicellular spheroids and large multicellular masses as three distinct, previously unrecognized DTP states that expand the spectrum of drug-tolerant persistence. Notably, the large round cells continued to increase in size, although a subset underwent cell death during this process, resulting in total cell volumes approaching those characteristics of deep quiescence or low-cycling states. Additionally, a minority of daughter cells derived from multicellular structures survived, differentiated into small round cell intermediates, and transitioned back to a multicellular state (Fig. 3j). This reveals a cyclic route that preserves a minimal DTP “seed” population under prolonged PTX pressure (Fig. 3k). Therefore, it is possible that the DTP trajectory is not a random walk but a programmed, drug-directed differentiation cascade: the large round mononuclear cells → multicellular spheroids→ large multicellular masses (Fig. 3k). This stepwise progression indicates that PTX awakens an evolutionarily conserved developmental program rather than merely evoking stochastic transcriptional noise.

### Induced formation of oocyte-like state

Building on the observations that (i) a subpopulation of large round cells exhibits oocyte/blastomere-like morphology and (ii) we previously revealed that a subset of somatic tumor cells can re-enter a blastomere-like state through gametogenesis- and parthenogenesis-like programs, we investigated whether a germline/early embryonic differentiation trajectory fuels multicellularity. Alkaline phosphatase (AP), a marker of germ cells and blastomeres^8^, was used to track the sequential emergence of germ cell-and early-embryo-like cells following drug exposure. Consistent with the higher of big round-shaped cell formation, the number of AP positive oocyte-like cells abrupt increased in 4T1 culture treatment with PTX and VCR compared with that treatment with DDP, DOX and 5-FU for 24h (Fig. S3), suggesting that the formation of large round-shaped cells is possibly linked to oocyte-like cell generation. We then employed PTX as a model to dissect the germ cell- and early-embryo-like transition. Strikingly, within 4 hours of PTX (10 µM) exposure, AP-positive PGC-like cells began to exhibit increased diameter and cell numbers, eventually exceeding 20 µm in diameter, suggesting development toward oocyte-like cells; by 8 hours, oocyte-like cells had increased ∼100-fold versus vehicle-treated controls, and by ∼12 hours, a subset transitioned to weakly AP-positive or negative states (Figs. 4a-4e, Tables S12-14), indicating sequential progression resembling physiological oocyte formation. The number of oocyte-like cells peaked approximately 8 hours (comprising ∼40-50 % of all cells) after treatment with 10 µM PTX and subsequently declined (Figs. 4a-4e, Tables S12-14). Twenty-four hours post-treatment, ∼20 % of the cells displayed an oocyte-like phenotype (>20 µm in diameter) (Figs. 4a-4e, Tables S12-14).

**Fig. 4.**
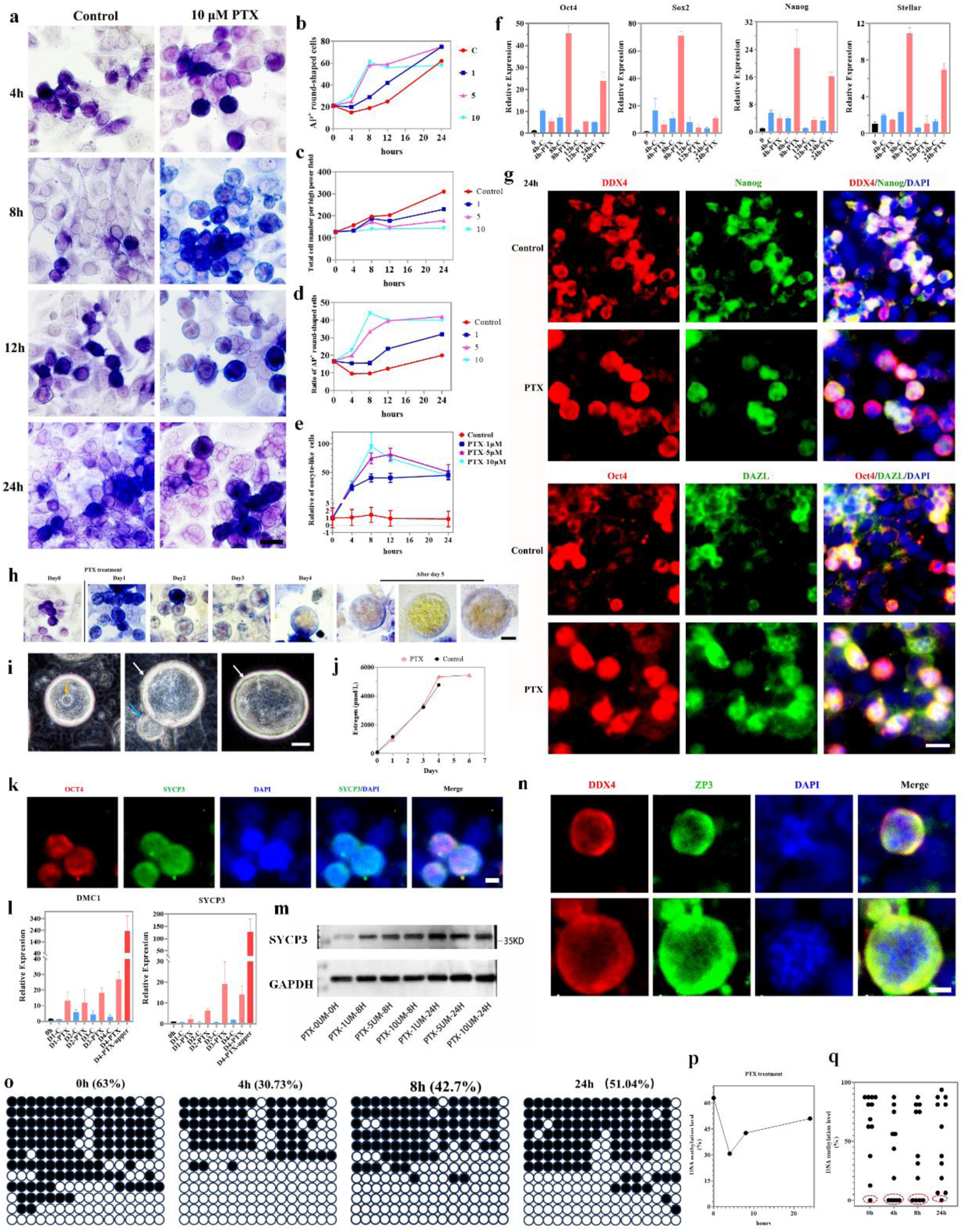
PTX induces oocyte-like cell formation in 4T1 cultures. (a) Bright-field images and alkaline phosphatase (AP) staining of control versus 10 µM PTX-treated cultures at sequential time points. (b–e) Quantification of AP⁺ cells, AP⁺ large round cells, and oocyte-like cell formation capacity (n=30). (f, l) RT-PCR analysis of germline and pluripotency gene expression at indicated time points (n=3). (g, k, n) Immunofluorescence staining of germline markers in PTX-treated cultures (24–48 h). (h) AP activity dynamics in germ-cell-like cells at distinct developmental stages. (i) Representative bright-field images of oocyte-like cells showing germinal vesicle (GV)-like nucleus (orange arrow), polar-body-like extrusion (blue arrow), and zona-pellucida-like rim (white arrow). (j) Estrogen concentration measurements in culture supernatants. (m) Western blot analysis of germline and meiotic proteins following PTX treatment. (o–q) Sodium bisulfite sequencing (o) and quantitative analysis (p, q) of H19 DMR methylation dynamics (● methylated; ○ unmethylated CpG sites). Scale bar = 25 μm (a, i), 10 μm (g, k, n). Statistical significance: *P < 0.05, **P < 0.01, ***P < 0.001, ****P < 0.0001.

Treatment with PTX for 8 and 24 hours significantly elevated the mRNA expression of Oct4, Sox2, Nanog, and Stella, as determined by RT-PCR, compared to untreated controls (Figs. 4f and S4a, Table S15). Consistent with this morphological switch, immunostaining revealed that numerous oocyte-like cells were positive for DDX4, Nanos3, DAZL, Oct4, Nanog and ZP3^8,9^ (Figs. 4g, S4b and S4c), supporting their similarity to natural oocytes. Immunofluorescence analysis further revealed a defined maturation trajectory: NANOS3 and Nanog (earlier marker) expression decreased whereas DDX4 (late marker) expression increased (Figs. 4g, S4b and S4c). These coordinated changes occurred in tandem with extended culture time and increasing cell diameter, indicating that the chemical treatment orchestrates a progressive fate transition from a PGC-like to an oocyte-like state. Collectively, these findings suggest that PTX stress promotes the acquisition of an oocyte-like identity in 4T1 cultures.

Of note, prolonged treatment allowed a subset of oocyte-like cells to further progress (Fig. 4h), generating markedly larger forms (∼50–100 µm) that detached or floated in the medium, consistent with a more mature phenotype. A subset of the oocyte-like cells acquired classic oocyte hallmarks: a prominent GV-like nucleus, polar-body-like extrusions and a translucent zona-pellucida-like rim (Fig. 4i). High level of estrogen could be detected in the 4T1 cultures, providing a support for the mature of oocyte-like mature (Fig.4j, Table S16). Both SYCP3 transcript and protein were detected in PTX-treated 4T1 cultures (Figs. 4k-m, Table S17), indicating that some germ cell-like cells had initiated meiotic entry. Karyotyping further captured meiotic figures in a subset of drug-treated cells—a configuration rarely, if ever, observed in controls (Fig. S4d). Additionally, these late-stage oocyte-like cells stained weakly or negatively for AP staining, while uniformly positive for DDX4 and ZP3 (Fig. 4n), resembling the immunophenotype of native oocytes. Collectively, these findings further indicate that the oocyte-like cells bear resemblance to natural oocytes with respect to advanced maturational characteristics.

Epigenetic signatures, most notably DNA methylation at germline differentially methylated regions (DMRs), are extensively and dynamically reprogrammed during normal female germ-cell development. To verify that PTX drives somatic cells toward an oocyte-like fate via a PGC-like intermediate, we monitored time-resolved changes in H19-DMR methylation. Sodium-bisulfite sequencing demonstrated biphasic H19-DMR methylation dynamics in 4T1 cells upon PTX treatment—initial imprint erasure (4 h) resembling primordial germ cell development was followed by locus-wide remethylation (24 h), recapitulating the remethylation wave accompanying female germ-line maturation (Fig. 4o-q). This sequential reprogramming provides evidence that PTX stress alone is sufficient to trigger phenotypic plasticity and rapid entry into an oocyte-like developmental trajectory.

### Induced the formation of blastomere-like state

It is well documented that oocytes can give rise to blastomeres via parthenogenesis, a reproductive strategy common in certain lower organisms^10^. In humans, however, the same process is pathological and gives rise to ovarian teratomas^9^. We subsequently investigated whether blastomere-like cells were really induced by PTX treatment. Following 24 h of exposure to 10 µM PTX, the numbers of blastomere-like cells increased ∼100-fold versus controls in a dose-dependent manner and were most frequently observed at the 2- to 8-cell stage (Figs. 5a-d, Table S18). The numbers of blastomere-like cells rose abruptly at ∼12 h and peaked at ∼24-48 h post-exposure to 10 µM PTX, lagging the oocyte-like peak by only a few hours (Figs. 4e and 5d, , Table S18), indicating blastomere-like progeny possibly via parthenogenetic activation of oocyte-like cells. Combined AP staining, HE staining, immunostaining for pluripotency markers (Oct4, Sox2 and Nanog) and oocyte-specific glycoprotein ZP3 revealed blastomere-like cell identity within PTX-induced 4T1 cultures (Figs. 5b, 5c, 5e, 5f, S5a and S5b). Notably, heterogeneous AP activity among multinuclear cells suggested that blastomere-like identity was restricted to a subpopulation (Figs. 5a and S5c). The findings indicate the rapid, stress-driven conversion of 4T1 cultures into blastomere-like progeny.

**Fig. 5.**
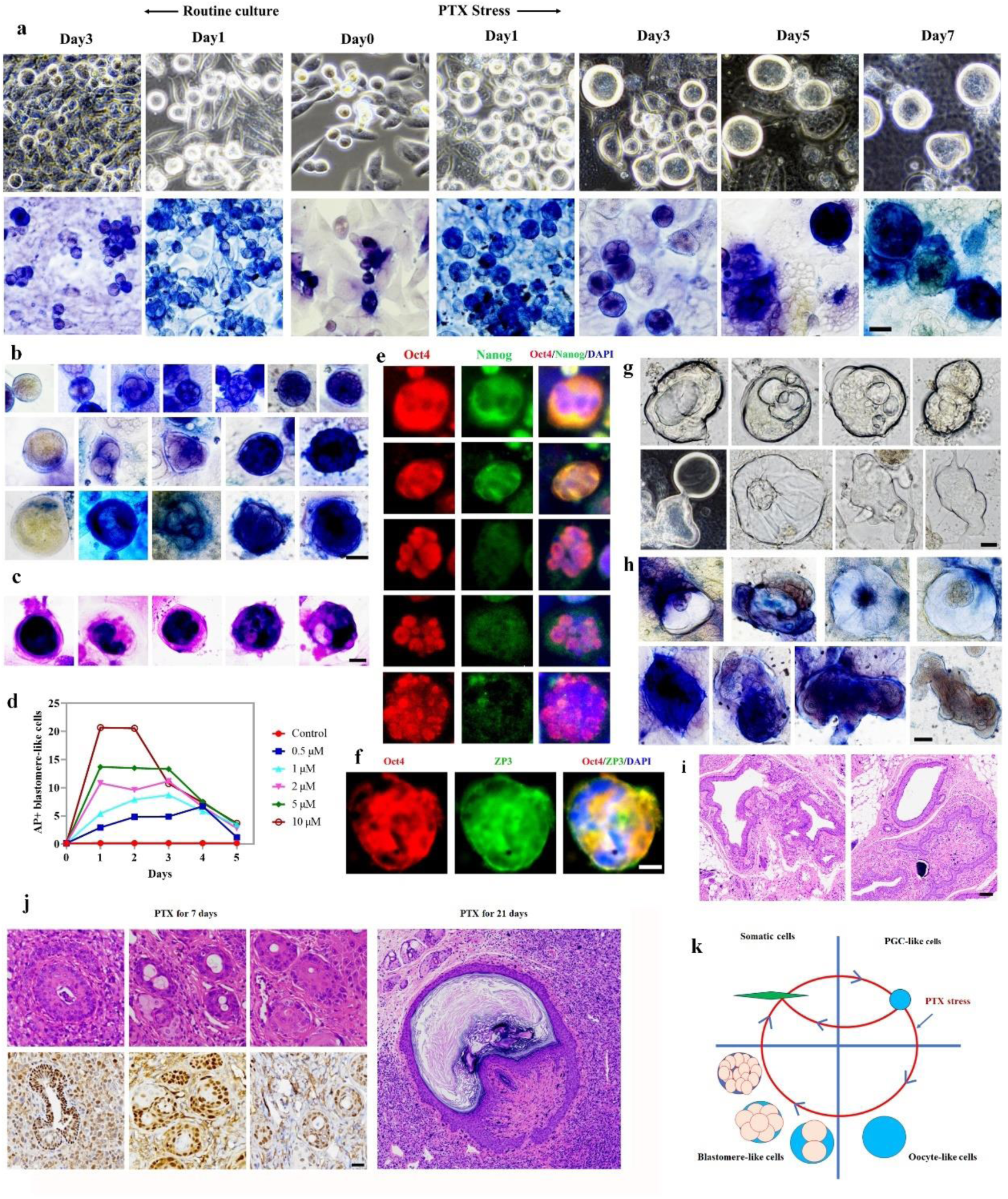
PTX induces blastomere-like cell formation in 4T1 cultures. (a) Bright-field images and AP staining of control versus 10 µM PTX-treated cultures at sequential time points. (b, c) AP and H&E staining showing progressive developmental stages from oocyte-like to blastomere-like cells. (d) Quantification of AP⁺ blastomere-like cells across PTX concentrations and time points (n=9). (e, f) Immunofluorescence of early embryonic markers in PTX-induced blastomere-like cells. (g, h) Bright-field and H&E imaging of multicellular masses. (i, j) H&E and immunohistochemical staining of tumor sections from suspended PTX-pretreated cultures (i, 48 h) and established 4T1 xenografts (j, 7/21 days PTX exposure). (k) Schematic model of blastomere-like cell trajectory under PTX stress. Scale bar = 25 μm (a–c, h), 10 µm (e, f), 40 µm (i, j). *P < 0.05, **P < 0.01, ***P < 0.001, ****P < 0.0001.

From days 1–3 following 10 µM PTX post-treatment, some blastomere-like cells stabilized at a similar diameter and, resembling native blastomeres, underwent successive cleavages without increasing in size; with each division, their volume was halved, resulting in exponential shrinkage to a densely compacted, size-compressed morphology (Fig. 5a and 5b). A subpopulation of blastomere-like cells released daughter cells that were positive for AP staining (Fig. S5d). As culture proceeded, blastomere-like cells segregated into three distinct size cohorts—∼25 µm, 26–50 µm and >50 µm—mirroring the heterogeneous maturation stages of their parental oocyte-like cells (Fig.5b). Prolonged culture induced TNAP/Oct4 downregulation and multicellular development (Fig. 5g, 5h). From ∼8–9 days post-treatment, cultures generated post-implantation embryo-like structures that ultimately formed large masses but collapsed into disorganized piles upon extended culture (Fig. 5g, 5h).

To further validate their similarity to native blastomeres, suspension cultures highly enriched (>95%) in blastomere-like cells and oocyte-like cells, generated by treating 4T1 cells with 10 µM PTX for 2 days, were injected into mice. Strikingly, these cultures could form teratomas (Fig. 5i), providing in-vivo functional evidence that a subset of blastomere-like cells persisted along an early-embryonic trajectory. Extending this observation, we first established 4T1 xenografts, allowed tumors to reach ∼60 mm³, and then challenged them with PTX. After 7 or 21 days of therapy, the explants had converted into tumors harboring teratomatous components—embryoid bodies and multiple differentiation (Fig. 5j), indicating that PTX triggers 4T1 cells toward an embryo-teratoma-like fate and multicellularity *in vivo*. Collectively, these studies revealed that PTX stress enable induce 4T1 cancer cells from a unicellular state switch to a blastomere-like, multicellular state (Fig. 5k).

### Activation of PGC-like state links drug stress-induced multicellularity

The data presented above failed to resolve the lineage derivation of blastomere-like state, leaving ambiguous whether they stem from a PGC-like or somatic antecedent. To further investigate whether PGC-like state is essential for the transition, we exposed 168FARN, an isogenic, lower-malignancy derivative with a constitutively smaller PGC-like compartment^11^, to identical PTX stress. These cultures generated markedly fewer oocyte-like cells, blastomere-like cells and multicellular aggregates (Figs. 6a and 6b, Table S19), indicating that acquisition of a PGC-like identity is critical for the rapid unicellular-to-multicellular switch. Moreover, multicellular plasticity is not intrinsic to the cellular program but represents an evolutionarily acquired phenotype, selectively favored and retained during malignant progression to aggressive states.

**Fig. 6.**
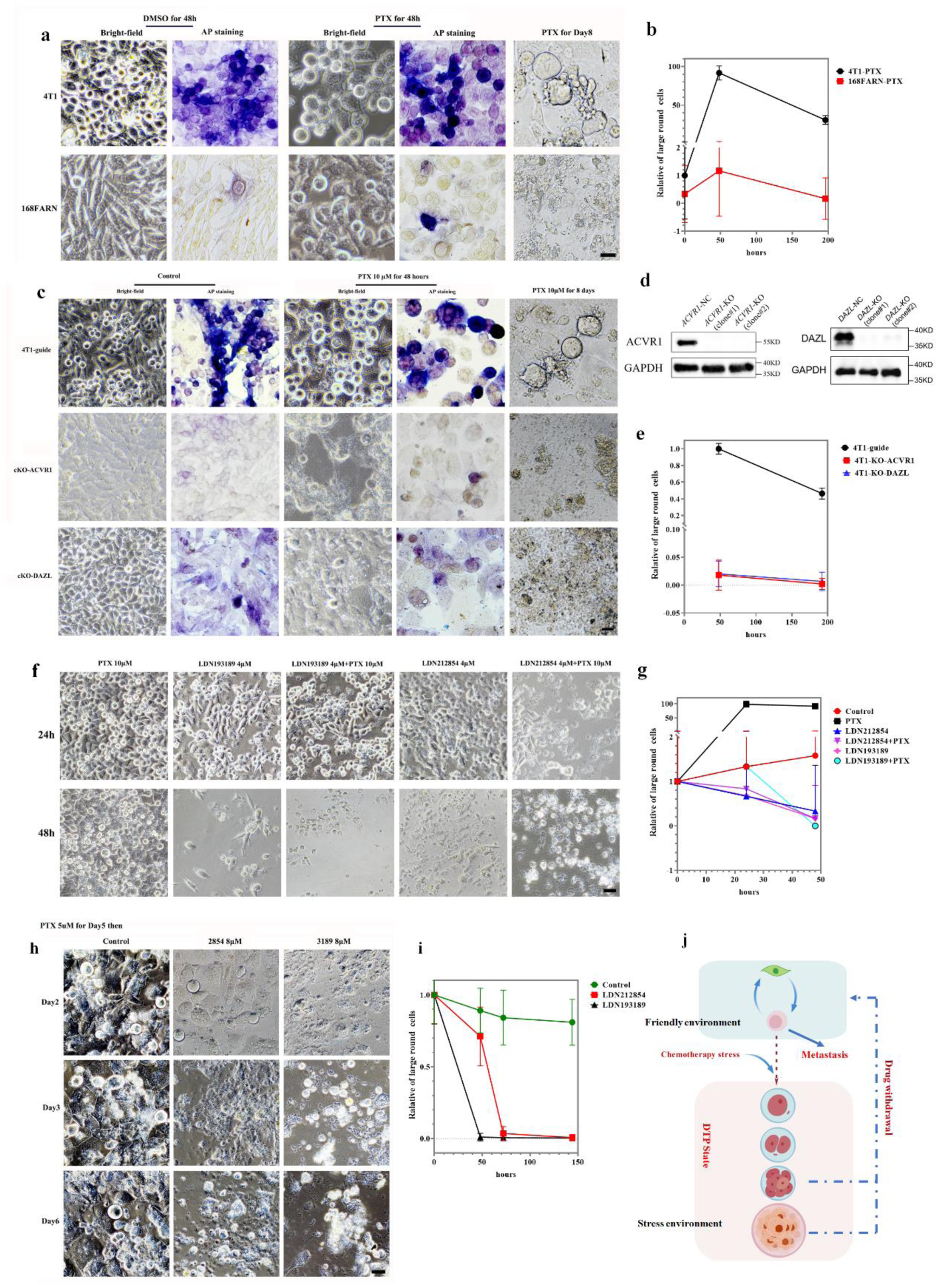
Essential role of PGC-like state in PTX-induced multicellularity. (a, b) Differential capacity for 10 µM PTX-induced large round-cell formation between 4T1 and 168FARN cultures. (c–e) Divergent formation capacity in control versus ACVR1 or DAZL knockout 4T1 cells (bright-field imaging, Western blot, and quantification). (f, g) Drug-induced large round-cell formation under indicated treatments. (h, i) Survival of large round cells from 5-day PTX-treated cultures after drug withdrawal and compound exposure. (j) Working model integrating new findings (yellow) with prior work connecting distinct malignant behaviors. Scale bar = 25 μm. *P < 0.05, **P < 0.01, ***P < 0.001, ****P < 0.0001.

To further confirm this concept, we then knocked out key genes associated with PGC specification, including *ACVR1* and *DAZL*, and then treated these knockout (KO) cells with PTX. The results showed that upon knockout of *ACVR1* or *DAZL*, 4T1 cells exhibited a significant reduction in the formation of PGC-like cells and a corresponding decrease in PTX-induced oocyte-like, blastomere-like cells and unicellular-multicellular transition (Figs. 6c-e, , Table S20). These data further suggest that both oocyte-like, blastomere-like cells and multicellular mass arise via a PGC-like intermediate state.

Given prior reports^8,12^ and our own work^11,13^ indicating that BMP signaling is essential for oocyte development, early embryogenesis, and SPLC formation, we asked whether pharmacologic blockade of the BMP receptor ACVR1 would suppress unicellular-multicellular transition and survival. In line with our earlier findings, 4T1 cells proved highly sensitive to the ACVR1 inhibitors LDN193189 and LDN212854; exposure to 8 µM of either compound for 24 h induced widespread cell death (Fig. 6f). To determine whether ACVR1 activity is required for PTX-induced blastomere-like cell formation, we co-treated 4T1 cells with 10 µM PTX plus 2 or 4 µM of LDN193189 or LDN212854 for 24 h. At 4 µM, either inhibitor completely abolished the emergence of unicellular-multicellular transition; by 48 h, virtually all cells had succumbed to the inhibitors (Figs. 6f and 6g, , Table S21). Next, we investigated whether established blastomere-like cells remained dependent on ACVR1 signaling. Cultures that had been primed with 10 µM PTX for 5 days to generate oocyte-like cells and multicellular cells were challenged with 8 µM LDN193189 or LDN212854. Within 48 h, the inhibitors eradicated virtually all identifiable cell subpopulations, including the oocyte- and blastomere-like fractions, and no viable cells remained after 5 days (Figs. 6h and 6i, Table S22). Collectively, these data indicate that ACVR1 not only licenses the formation of multicellular structures but also sustains their survival and chemo-resistance. Pharmacological blockade of ACVR1 with LDN193189 or LDN212854 thus represents a promising therapeutic strategy against 4T1-like malignancies.

## Discussion

In this study, we reveal that certain cancer cells undergo unicellular–multicellular bistable plasticity in response to PTX stress and withdrawal via a non-genetic mechanism, thereby enhancing fitness under chemotherapeutic pressure and generating a distinct DTP population. Notably, this transition is blocked when PGC-specific genes are silenced and the inhibitor itself impairs PGC specification. Collectively, our findings enhance our understanding of cancer cell plasticity, evolutionary trajectories (direction), phenotypic adaptation, and the emergence of DTP states.

Traditionally regarded as traitors to multicellularity, cancers are thought to abandon cooperative constraints and revert to a unicellular lifestyle to maximize autonomy and fitness^14–16^. Conversely, across the tree of life—eukaryotes, bacteria, archaea, haloarchaea and beyond—environmental stress repeatedly drives single cells to assemble into multicellular forms such as biofilms or clonal aggregation, strategies that enhance stress tolerance and survival^3,4,17^. Our data reveal that chemotherapeutic pressure enables cancer unicells to retrace the evolutionary transition to clonal multicellularity^18,19^. Rather than a linear progression, neoplastic history may therefore be circular: multicellularity → unicellularity → stress-induced multicellularity, mirroring the universal rule that stress favors collective survival and might act as a key engine of tumor evolution^20,21^. This reversible trajectory offers an experimentally tractable model for dissecting the evolutionary transition from single cells to multicellular organisms^14^ and simultaneously indicates that cancer’s evolutionary arrow points toward an independent, quasi-autonomous life form.

DTP cells are the chief engines of post-treatment relapse, yet their survival strategies and evolutionary trajectories remain unclear^1,2,22^. Current catalogs scatter them across cancer-stem, senescence-like arrest, quiescence, lineage plasticity, diapause and onco-fetal reprogramming^2,23–25^, invoking either stochastic or deterministic fate transitions; however, classical stem markers (such as OCT4, SSEA1, CD117) overlap with primordial-germ-cell (PGC) signatures, and both diapause and onco-fetal modules are inherently embryonic^26^, implying that a sizeable fraction of previously described DTPs already map onto a germ-line/embryonic axis. Our study reveals a new DTP state and a longitudinal framework that reframes DTPs as fluid phenotypic trajectories rather than fixed subclasses ^27^, providing direct evidence for adaptive evolution in cancer cells.

This aligns with an expanding body of evidence that stress can drive cancer evolution before mutations pile up—a Lamarckian flash in a Darwinian world^28^. Still, the ability to make that shift is mutation-built: earlier DNA hits confer the epigenetic potential that permits the lifestyle switch. Consequently, only cell lineages that have accumulated the requisite mutational set can multicellularity, explaining why identical drug stress induces weak or incomplete reprogramming in other clones. Cancer evolution is therefore neither purely Darwinian nor purely Lamarckian^28–31^, but a sequential, two-step duet: Darwinian mutation and selection build the epigenetic tool-kit while Lamarckian induction rapidly deploys that tool-kit for immediate adaptation. Importantly, this epigenetic-driven, reversible transitions supply the fastest response to hostile niches and survival, buying time through repeated transitions until a genetic mutation that stably fits the hostile niche is ultimately acquired.

In addition, it was well documented that polyploid giant cancer cells (PGCCs), notably large and multinucleated, frequently emerge in advanced tumors or after therapy^32,33^ display blastomere-like traits and cause DTP state and cancer relapse ^34,35^, therefore a subpopulation of PGCCs may represent the most visible tip of multicellular reprogramming. Our findings extend beyond morphological description^33^, revealing not only the generation of PGCCs, but also their subsequent progression into multicellular masses and the establishment of environment-dependent unicellular–multicellular bistable plasticity.

In metazoans, the transition to clonal multicellularity proceeds from embryonic stem cells through primordial germ cells and oocytes to blastomeres. Tumorigenesis mimics gametogenesis and early embryogenesis at both molecular and organizational levels^26,36–38^, prompting Old to propose that cancer is a re-activated gametogenic program—a “disease of reproduction”^36^ or “somatic-cell pregnancy”^39^. We previously identified two programs in tumor cells ^9,13,40,41^: a “minor cycle” (somatic → ESC/PGC-like → somatic) and a “major cycle” (somatic → ESC/PGC-like → oocyte-like → parthenogenetic blastomere-like → somatic). The present study reveals that cytotoxic stress triggers stepwise progression along the embryonic/germ-cell axis, driving entry into the major cycle and a DTP state (Fig. 6j). Conversely, a supportive microenvironment activates the minor cycle, fueling explosive proliferation. Environmental stimuli therefore evoke bistable plasticity, dynamically alternating between these cycles to maximize clonal fitness (Fig. 6j). Strikingly, the literature and our previous work have suggested that embryonic/germ-cell traits are central drivers of cancer malignancy^9,13,36–38,41–46^. Together, the findings lend strong experimental support to Old’s concept^36,39^ and our novel gametogenesis hypothesis of tumors^9,40^ and underscore that phenotypic plasticity—rather than discrete subclonal lineages—constitutes a dynamic continuum in which position along the embryonic/germline axis coordinates core neoplastic phenotypes (such as metastasis and DTP state) and intratumoral heterogeneity (Fig. 6j).

### Limitations of study

However, further investigation is needed to clarify the crucial role of the blastomere-like state in multicellularity and the molecular mechanisms underlying stress-induced adaptive evolution toward multicellularity in cancer and the key genetic mutations that drive the completion of this evolutionary process. In addition, the key genetic alterations required for the completion of this evolutionary transition.

## Acknowledgments

We acknowledge the use of AI-assisted language editing tools to improve grammar and writing clarity. These tools were employed solely for linguistic refinement, and all scientific content, data interpretation, and conclusions remain the original work and responsibility of the authors. This work was supported by the National Natural Science Foundation of China (No. 82273113) to C.L., National Natural Science Foundation of China (No. 82372327) to Z. M.

## Author contributions

J. L., Z. M., Z. Z., and E.Z. performed the most of experiments. Z. M. draw a part of cartoon and designed part of the experiments. C. L. designed the experiments, analysed the data and wrote the manuscript.

## Availability of supporting data

All data are available in the main text or in the supplementary materials.

## Consent for publication

Not applicable

