## Supplemental materials for "Chemotherapy-induced multicellularity drives drug-tolerant persistence state in tumor cells"

### STAR METHODS

### EXPERIMENTAL MODEL AND STUDY PARTICIPANT DETAILS

#### Cell Lines

The 168FARN cell line (obtained from Dr. Kounosuke Watabe) and the 4T1 cell line (from ATCC) were both established from the same breast tumors grown in BALB/c mice. The human cell lines HEK293T were acquired from the Cell Bank of the Chinese Academy of Science (Shanghai, China). The human HEK293T cell line was obtained from the Cell Bank of the Chinese Academy of Sciences (Shanghai, China). All cells were maintained in Dulbecco's Modified Eagle Medium (DMEM; Gibco) supplemented with 10% fetal bovine serum (FBS; Sigma-Aldrich), 100 units/mL penicillin, and 100 µg/mL streptomycin (Gibco), and incubated at 37°C in a humidified incubator with 5% $CO_2$. Sex information for these cell lines was not specified. No additional cell line was utilized in this investigation.

#### Animal Models

Female BALB/c nude mice (4–6 weeks old) were purchased from Shanghai Model Organisms. The mice were maintained under standard housing conditions at controlled temperature with free access to food and water under a natural light/dark cycle. All animal procedures were reviewed and approved by the Animal Ethics Committee of Fudan University (Approval No. 2022JS Huashan Hospital-075) and conducted in accordance with the *Guide for the Care and Use of Laboratory Animals* (8th edition, National Academies Press).

### METHOD DETAILS

**Cell Culture**

All cells were cultured as described in the 'Cell lines' subsection above. For experimental assays, cells were seeded into multi-well plates during the logarithmic grow phase.

**Drug treatment**

4T1 cells were subjected to continuous exposure to DMSO or PTX (0.5, 1, 2, 5, or 10 μM), or treated with VCR, DDP, DXO, or 5-FU (each at 10 μM) for varying durations. The treated cells underwent microscopic imaging, AP staining, western blotting, immunofluorescence staining, animal experiments, karyotype analysis, or RT-PCR analysis.

**Replating cultures and generating daughter cells**

Spheres derived from 4T1 cells following various treatment durations were harvested, dissociated into single cells, and replated in fresh dishes overnight. The medium was subsequently removed, and cells were maintained in standard culture medium until daughter cells emerged.

**Drug withdrawal and generating daughter cells**

Drug-treated cells were washed to remove residual drug and subsequently cultured in drug-free medium. Half of the medium was replaced every 4 days until daughter cells appeared.

**Quantitative Real-Time PCR**

Total RNA was isolated using TRIzol reagent (Invitrogen) and reverse-transcribed into complementary DNA (cDNA) using the PrimeScript RT kit (Takara). Quantitative real-time PCR (qPCR) was performed on a 7500 Real-Time PCR System (Applied Biosystems) with SYBR Green Master Mix (Takara). Each reaction was performed in triplicate. The primer sequences are listed in Supplementary Table S23.

**Plasmids Transfection and CRISPR/Cas9 Gene Editing**

To disrupt the indicated genes, CRISPR/Cas9 (Addgene, #48138) and mouse sgRNA-RFP (Sigma) plasmids were utilized, with the guide empty vector (Sigma) acting as the control. For lentiviral particle generation, 20 µL X-tremeGENE9 DNA transfection reagent (Roche) was first diluted in 500 µL Opti-MEM (Gibco) and permitted to stand for 5 min. This solution was then combined with 6 µg lentiviral packaging plasmid (Beyotime, L00002M) plus 6 µg CRISPR/Cas9 plasmid for a 25-min incubation at room temperature. These mixtures were subsequently introduced into HEK293T cells suspended in 9.5 mL Opti-MEM medium. The medium was subsequently exchanged for DMEM supplemented with 10% fetal bovine serum (FBS; Sigma). The viral-containing medium from HEK293T cells was harvested following 24 h and 48 h of culture, filtered through a 0.45 µm Steri-Flip filter (Millipore), combined with 20% fresh culture medium containing 7 µg/mL polybrene, and applied to transduce 4T1 cells for 4 h. Following two rounds of transduction and 48 h of culture in standard 10% fresh medium, 4T1 cells were subjected to 4 µg/mL blasticidin for 5 days to generate 4T1-Cas9 cells. The indicated sgRNAs (Sigma) or guide plasmids (Sigma) were subsequently delivered into 4T1-Cas9 cells using the identical protocol. Cells were then exposed to 5 µg/mL puromycin for 5 days, after which single clones were isolated by limiting dilution in 96-well plates. Knockout cell clones were identified through DNA sequencing and western blotting. Sequencing primers, knockout cell sequencing results, and sgRNA information are provided in Tables S24-26, as well as our previous study.

**Western Blotting**

Protein extraction was performed using RIPA buffer supplemented with protease inhibitors. Extracts were resolved by SDS-PAGE and electroblotted onto PVDF membranes. Following blocking, the membranes underwent incubation with primary antibodies targeting DAZL (Novus Biologicals, NB100-2437), ACVR1 (Abcam, ab155981), and GAPDH (Gnimission, GNJ4110-GH). Signal detection employed HRP-conjugated secondary antibodies together with a chemiluminescent substrate.

**Immunofluorescence**

Cells cultured on chamber slides underwent fixation with 4% paraformaldehyde for 20 min at ambient temperature. Subsequently, samples were subjected to overnight incubation at 4°C with primary antibodies directed against Oct4 (1:400), Sox2 (1:200), DAZL (1:300), DDX4 (1:150), Nanog (1:200), Nanos3 (1:200), SYCP3 (1:200), and ZP3 (1:200). Following rinse steps, fluorescent-conjugated secondary antibodies were applied, specifically Goat Anti-Rabbit IgM/AF488 (Bioss, bs-0369M-AF488) and Goat Anti-Mouse IgG Fc/AF555 (Bioss, bs-0377R-AF555). Nuclear counterstaining utilized DAPI (Invitrogen).

**Alkaline phosphatase staining**

Cells were fixed with cold methanol for 7 min, followed by two rinses with Tris-HCl (pH 8.6). AP staining was subsequently performed using the Alkaline Phosphatase staining kit (Vector Laboratories) for 1 h, according to the manufacturer's protocol. The staining solution was then removed, and PBS was added to terminate the reaction. The stained cells were subsequently examined by microscopy.

**Animal Studies and Xenograft Model**

Female BALB/c mice (4–6 weeks old) were purchased from Beijing Vital River Laboratory Animal Technology Co., Ltd. All animal experimental protocols were approved by the Animal Ethics Committee of Fudan University (approval number: 2022JS Huashan Hospital-075) and performed in accordance with the Guide for the Care and Use of Laboratory Animals (8th Edition, 5th Revision, National Academies Press, USA). Mice were housed in an SPF animal facility at 22–24 °C with 40%–60% humidity and a 12 h light/dark cycle, with free access to standard chow and autoclaved water. Cells were inoculated subcutaneously at a density of $5\times10^6$ cells per mouse.

In the in vivo tumorigenicity assay, paclitaxel-induced sphere-derived daughter cells (4T1-TS5, 4T1-TS6), drug-surviving daughter cells (4T1-TA3), and drug-treated cells (4T1-PTX-D2, 4T1-PTX-D14) were inoculated separately, with untreated 4T1 cells serving as the control.

For the intratumoral administration experiment, 4T1 xenograft tumor models were first established. When tumor volume reached approximately 60 $mm^3$, tumor-bearing mice received intratumoral injection of paclitaxel at a dose of 15 mg/kg for 7 or 21 consecutive days. Control mice were injected with an equal volume of solvent simultaneously. The solvent consisted of 2% DMSO, 40% PEG400, 5% Tween-80, and 53% saline.

Tumor onset time was recorded and tumor-free survival curves were plotted during the experiment. Once visible tumors formed, the long diameter (L) and short diameter (W) of tumors were measured using an electronic digital caliper. Tumor volume was calculated using the formula $V = 1/2\times L\times W^2$, and tumor growth curves were generated. At the end of the experiment, mice were euthanized by carbon dioxide inhalation, and tumor tissues were collected and fixed for subsequent analyses.

**Histology and HE Staining**

For cultured tumor cells: fixation was performed in 10% neutral-buffered formalin (NBF) for 30 min, followed by hematoxylin and eosin (H&E) staining according to standard histological procedures. For tumor tissue samples: specimens were fixed in 10% NBF overnight, paraffin-embedded, and sectioned. Subsequently, sections underwent deparaffinization, rehydration, and H&E staining per routine histological protocols.

**Immunohistochemistry**

Antigen retrieval was conducted on deparaffinized sections using sodium citrate buffer (Biotend, RE202004) with microwave irradiation. This was followed by quenching endogenous peroxidase activity with 3% $H_2O_2$ and blocking non-specific binding sites with 5% BSA. Overnight incubation at 4°C employed primary antibodies targeting OCT4 (Abcam, ab184665) and SOX2 (Abcam, ab171380). Signal amplification utilized the Immunohistochemistry (IHC) Detect Kit for Rabbit/Mouse Primary Antibody (Proteintech, PK10006) together with DAB chromogen. Nuclear counterstaining applied hematoxylin (Solarbio, G1004), and coverslipping employed neutral balsam (Beyotime, C0173). For negative controls, PBS replaced primary antibodies.

**Quantification and statistical analysis**

Data are expressed as mean ± standard error of the mean (SEM) derived from a minimum of three independent replicates. All statistical comparisons were performed in GraphPad Prism 9. Two-group comparisons used unpaired t-tests, and multiple groups were compared by one-way ANOVA. Survival curves were generated by the Kaplan–Meier method and compared with the log-rank test. A p-value $< 0.05$ was considered statistically significant. Detailed sample sizes (biological replicates, animals) and specific statistical tests are indicated in the respective figure legends.

### ADDITIONAL RESOURCES

No additional resources (e.g., websites, forums) were generated or used in this study.

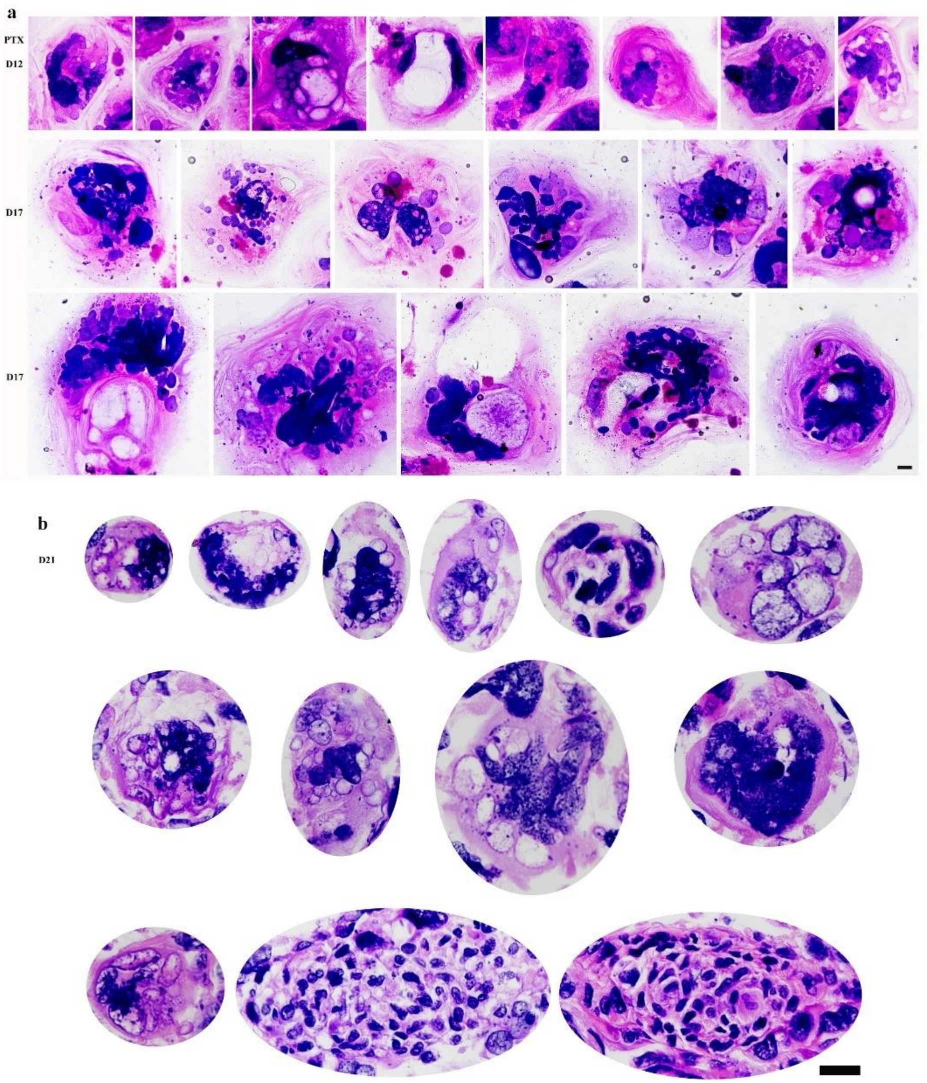
a
PTX
D12
D17
D17
b
D21

Supplementary Fig. S1. Bright-field microscopy of hematoxylin and eosin (HE)-stained 4T1 cell cultures treated with 10 μM paclitaxel (PTX). (a) Direct exposure to PTX. (b) Multicellular mass section after 21 days of treatment. Scale bar = 25 μm.

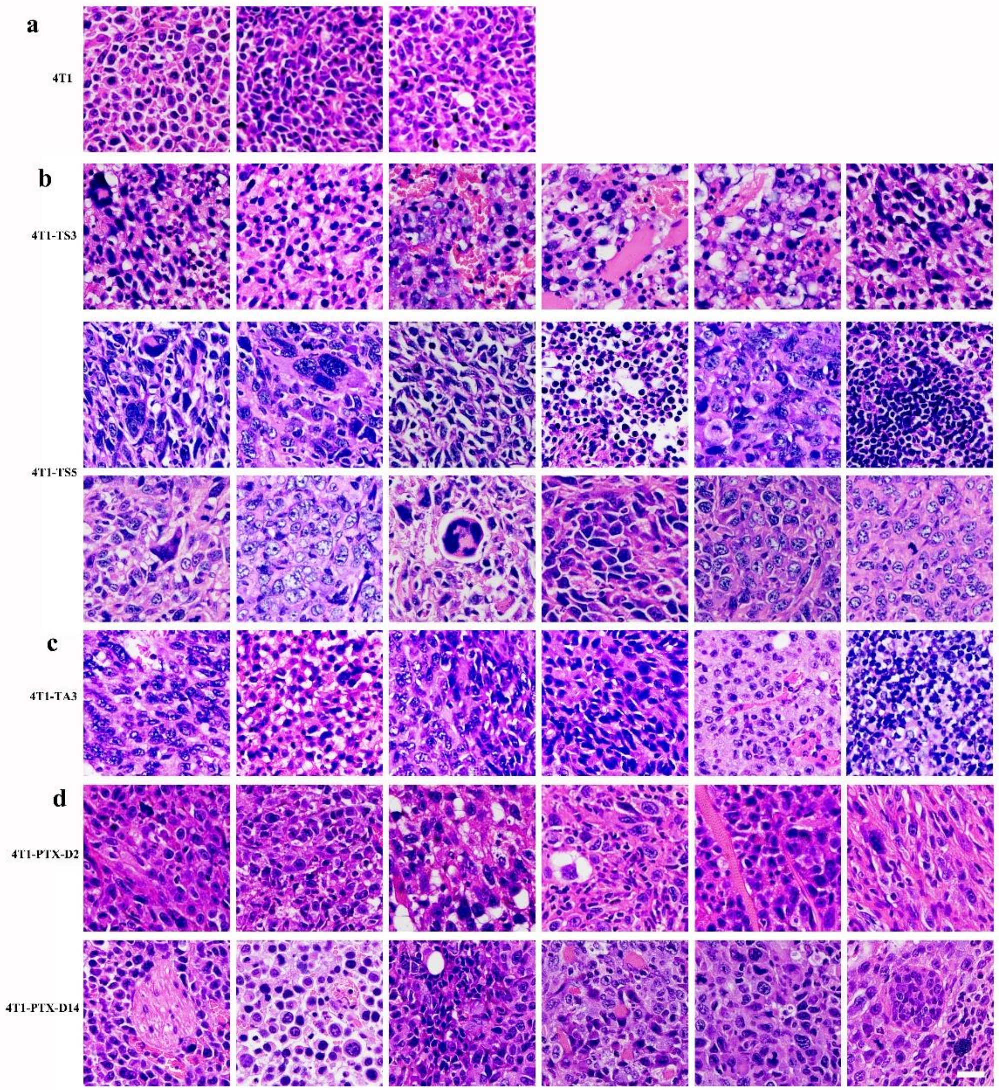
a
4T1
b
4T1-TS3
4T1-TS5
c
4T1-TA3
d
4T1-PTX-D2
4T1-PTX-D14

Supplementary Fig. S2 HE-stained tumor sections from 4T1-derived xenografts. (a) Untreated control. (b) Daughter cells from suspending spheres of drug-surviving cultures. (c) Daughter cells from attached drug-surviving cultures. (d) PTX-pretreated cells. Scale bar = 50 μm.

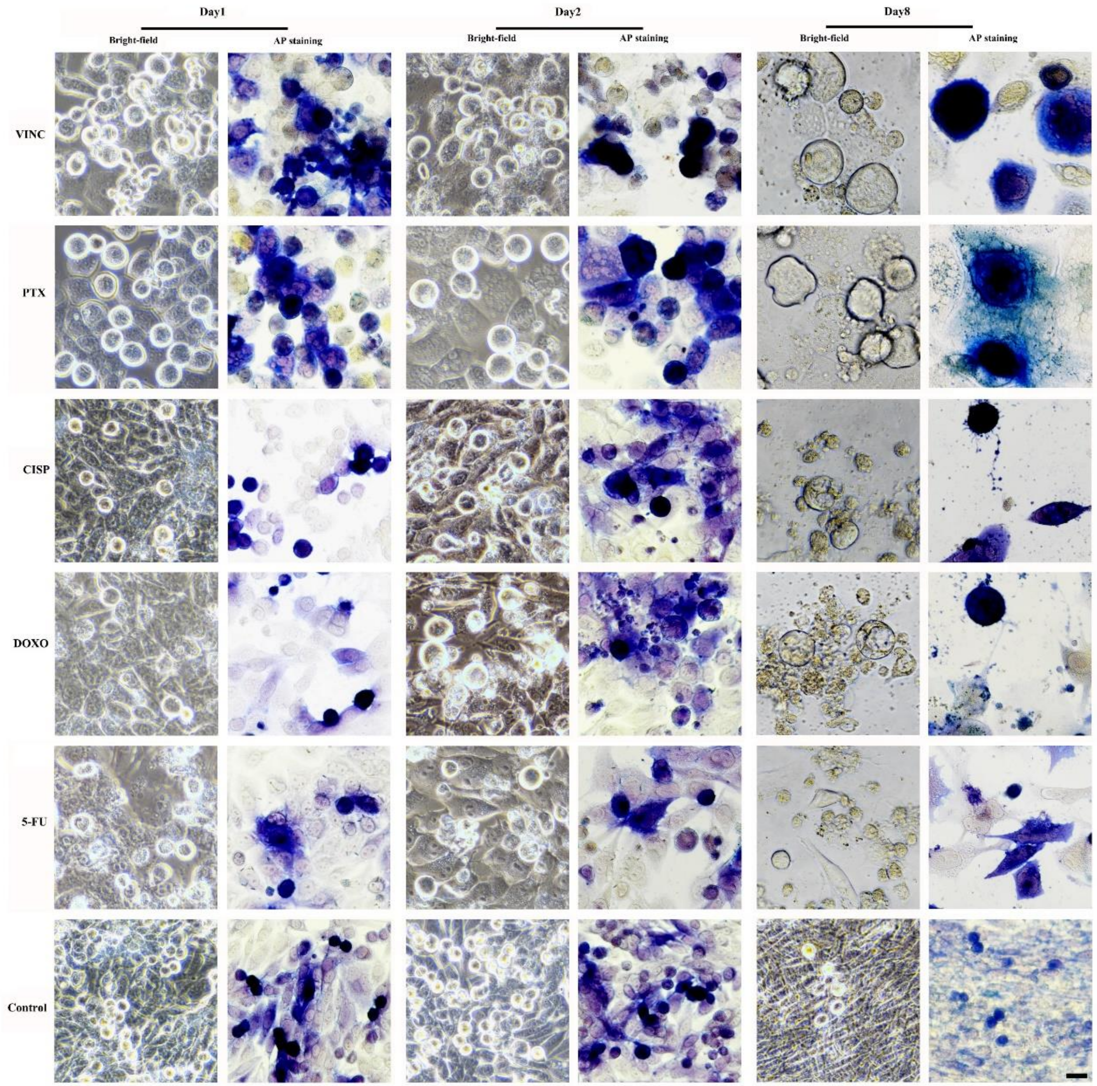

Day1
Day2
Day8
Bright-field
AP staining
Bright-field
AP staining
Bright-field
AP staining
VINC
PTX
CISP
DOXO
5-FU
Control

Supplementary Fig. S3 Characterization of alkaline phosphatase (AP)-positive large cells in 4T1 cells following chemotherapy treatment. (a) Representative phase-contrast and AP staining images at indicated time points across different chemotherapeutic agents and controls. (b) Quantification of AP-positive large cells across treatment groups (n=9 per group). Scale bar = 25 μm.

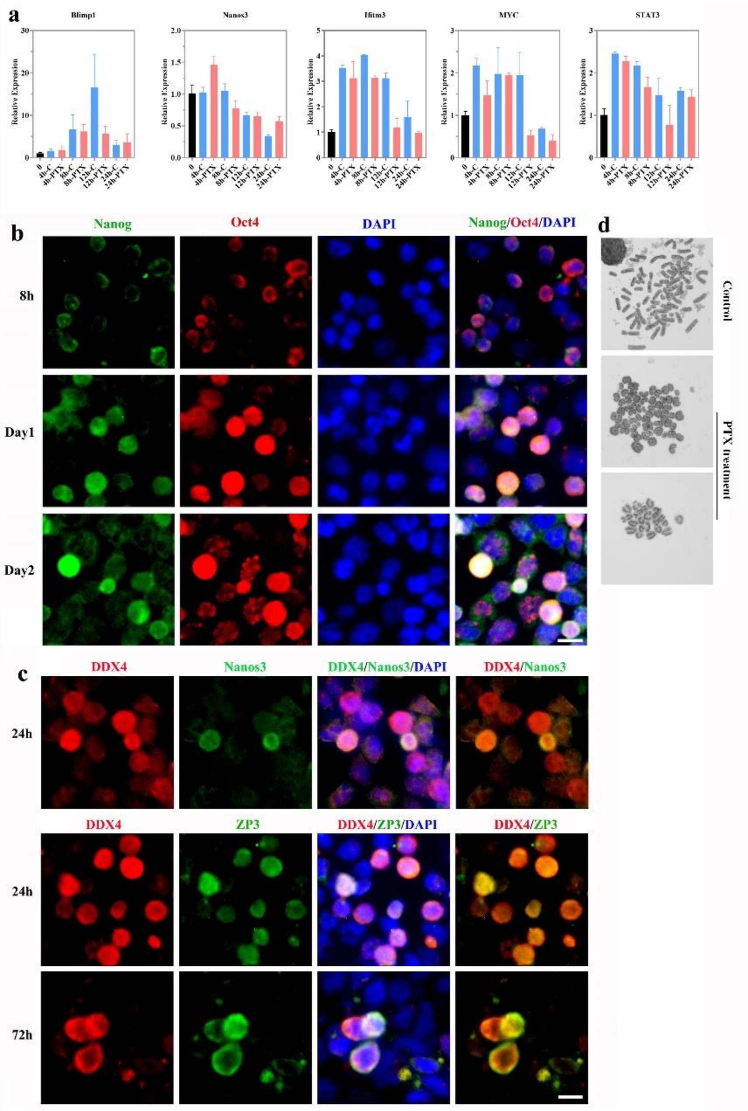
a
Blimp1
Nanos3
Ifitm3
MYC
STAT3
Relative Expression
b
Nanog
Oct4
DAPI
Nanog/Oct4/DAPI
8h
Day1
Day2
c
DDX4
Nanos3
DDX4/Nanos3/DAPI
DDX4/Nanos3
24h
ZP3
DDX4/ZP3/DAPI
DDX4/ZP3
72h
d
Control
PTX treatment

Supplementary Fig. S4 PTX treatment triggers oocyte-like cell differentiation in 4T1 cultures. (a) RT-PCR detection of germline-associated and pluripotency-related gene transcripts at sequential time points (n=3, per group). (b, c) Immunofluorescence imaging of germline markers following 24–72 h PTX exposure. (d) Karyotypic characterization of chromosomal integrity in control versus PTX-treated cultures. Scale bar = 25 μm.

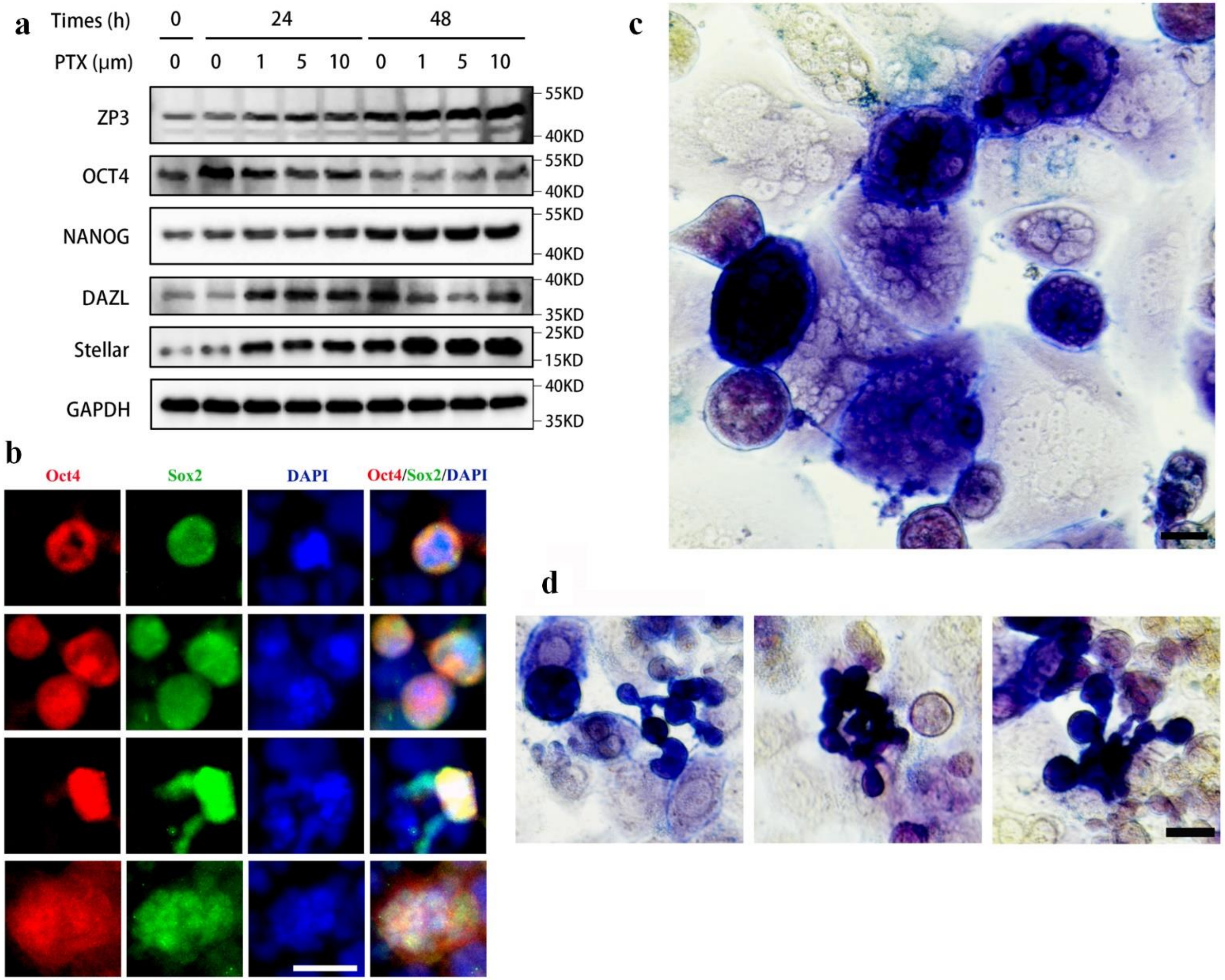

a
Times (h)
0
24
48
PTX (μm)
0 0 1 5 10 0 1 5 10
ZP3
OCT4
NANOG
DAZL
Stellar
GAPDH
55KD
40KD
55KD
40KD
55KD
40KD
40KD
35KD
25KD
15KD
40KD
35KD
b
Oct4
Sox2
DAPI
Oct4/Sox2/DAPI
c
d

**Supplementary Fig. S5. PTX induces blastomere-like cell formation in 4T1 cultures.** (a) Weston blot showed the expression of germline-associated and pluripotency-related protein at sequential time points. (b) Immunofluorescence imaging of pluripotency-related markers following PTX exposure. (c) AP staining showed distinct characteristics of the larger tumor cells. (d) AP staining showed the daughter cells from blastomere-like cells after treatment with PTX for 48 hours. Scale bar = 25 μm.

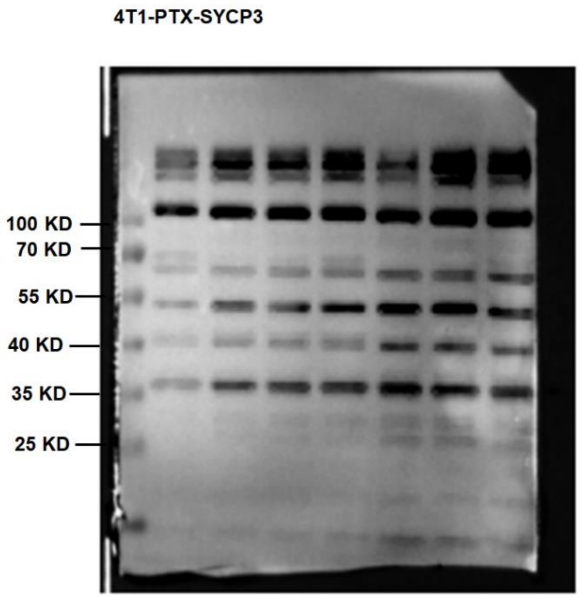

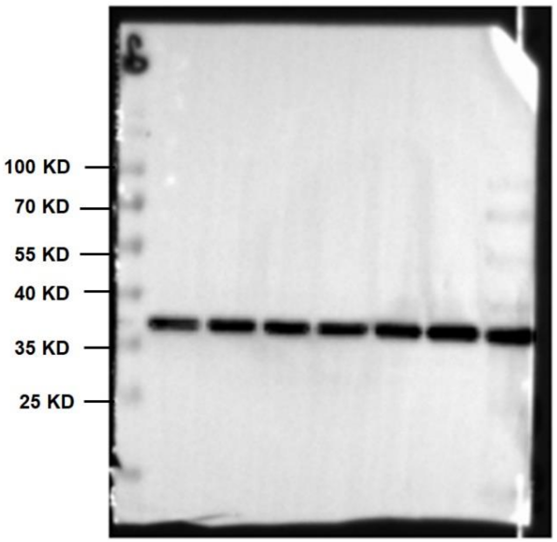

4T1-DAZL

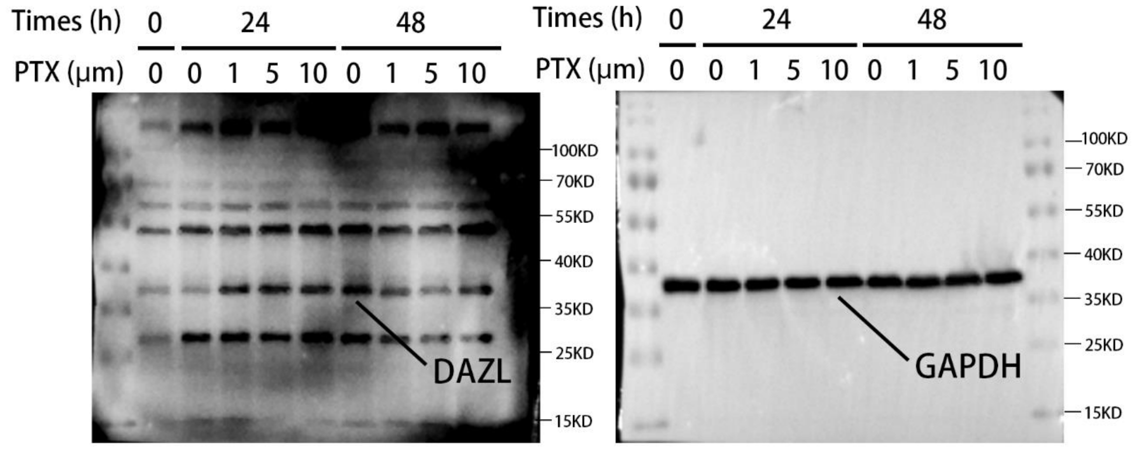

4T1-NANOG

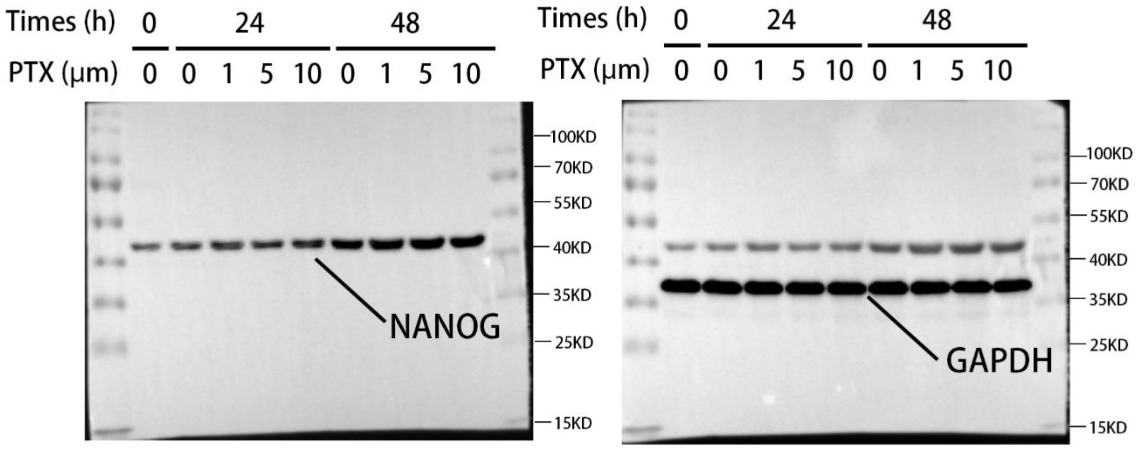

4T1-OCT4

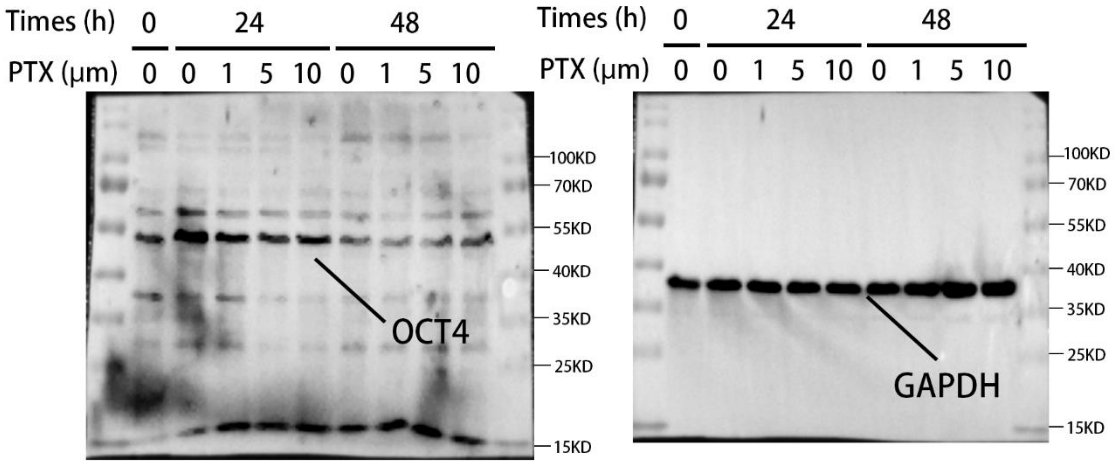

4T1-Stellar

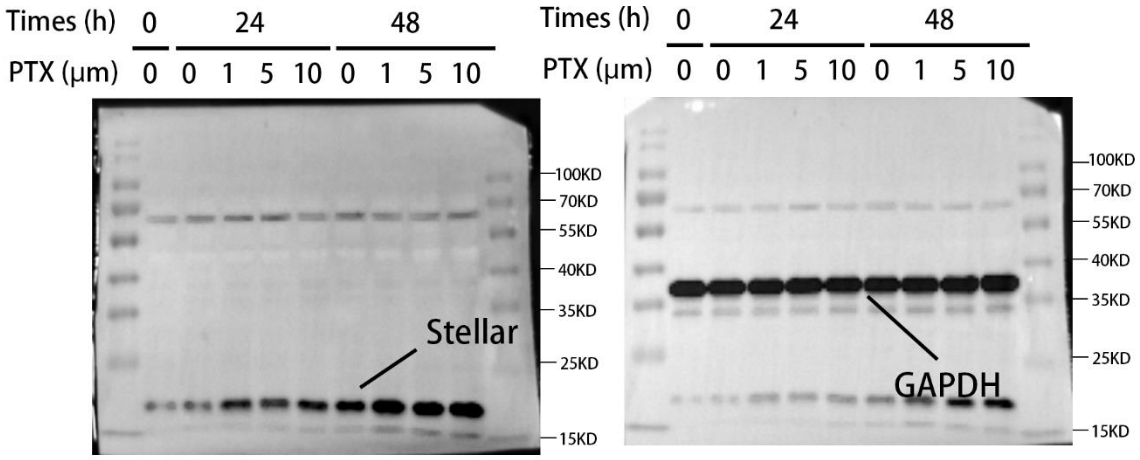

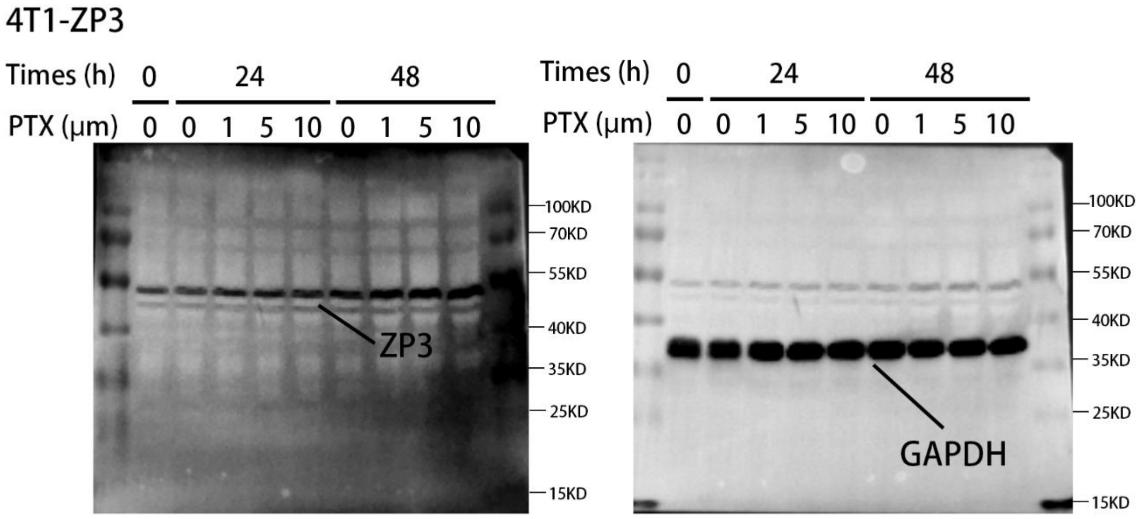
4T1-ZP3
Times (h) 0 24 48
PTX (μm) 0 0 1 5 10 0 1 5 10
ZP3
GAPDH
100KD
70KD
55KD
40KD
35KD
25KD
15KD

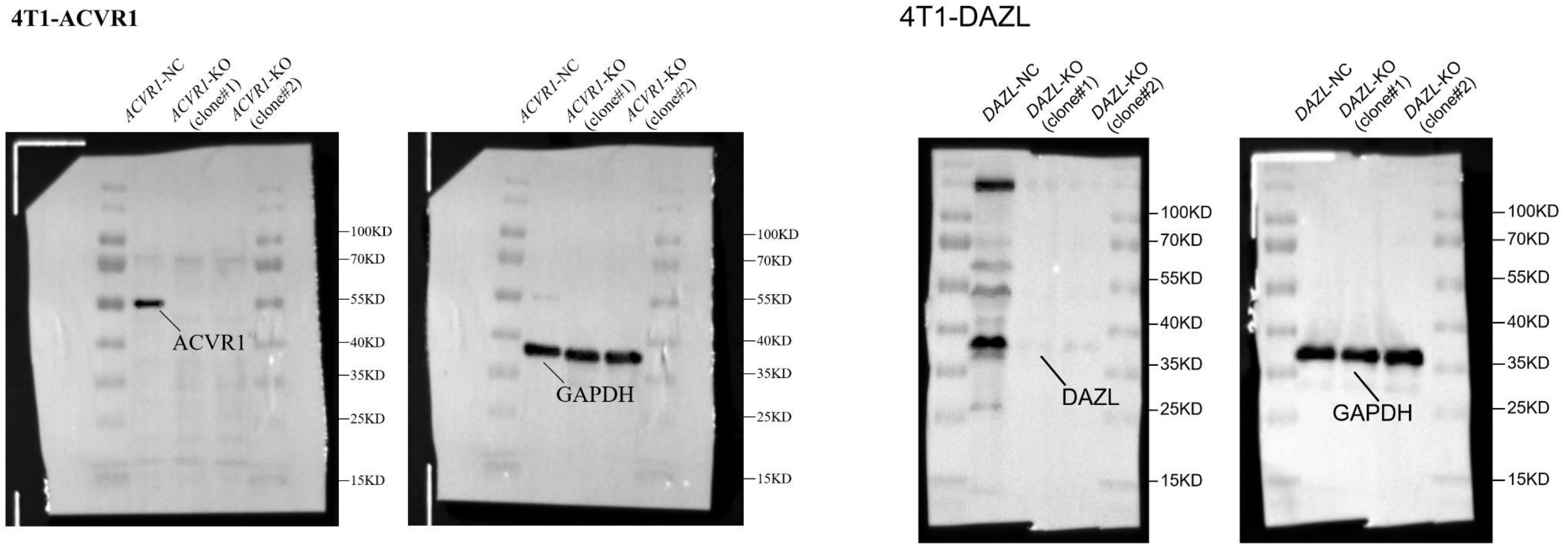

**Supplementary Fig. S6.** Images of the original western blot.

**Supplementary-Movie-1.** Morphological tracking of 4T1 cells during standard culture (days 0–4).

**Supplementary-Movie-2.** Morphological tracking of 4T1 cells during PTX treatment (days 0–4).

**Supplementary-Movie-3.** Morphological tracking of 4T1 cells during PTX treatment (6–9 days).

**Supplementary-Movie-4.** Morphological tracking of 4T1 cells during PTX treatment (11–14 days).

**Supplementary Tables**

**Table S1.** Large round-cell formation capacity per high-power field: comparison between control and drug-treated groups (n=20).

| | | Control | VCR | PTX | DDP | DOX | 5-FU |
|---|---|---|---|---|---|---|---|
| 0 | Mean | 0.25 | 0.25 | 0.25 | 0.25 | 0.25 | 0.25 |
| | SD | 0.55 | 0.001 | 0.11 | 0.07 | 0.35 | 0.12 |
| Day1 | Mean | 0.45 | 11.55 | 21.15 | 3.65 | 2.90 | 2.35 |
| | SD | 0.76 | 9.20 | 1.79 | 0.34 | 1.65 | 0.18 |
| | Control& | P value | <0.0001 | <0.0001 | <0.0001 | <0.0001 | <0.0001 |
| | | Summary | **** | **** | **** | **** | **** |
| Day2 | Mean | 0.55 | 11.53 | 18.05 | 6.60 | 7.75 | 4.55 |
| | SD | 0.66 | 6.98 | 0.70 | 0.54 | 1.88 | 0.43 |
| | Control& | P value | <0.0001 | <0.0001 | <0.0001 | <0.0001 | <0.0001 |
| | | Summary | **** | **** | **** | **** | **** |
| Day4 | Mean | 0.45 | 5.88 | 9.25 | 5.65 | 3.70 | 2.50 |
| | SD | 0.67 | 3.38 | 0.44 | 0.44 | 1.87 | 0.35 |

| | | | | | | | |
|---|---|---|---|---|---|---|---|
| | Control& | P value | <0.000001 | <0.000001 | <0.000001 | <0.000001 | 0.000005 |
| | | Summary | **** | **** | **** | **** | **** |
| Day8 | Mean | 0.50 | 2.20 | 6.05 | 0.90 | 1.05 | 0.25 |
| | SD | 0.69 | 1.95 | 0.39 | 0.22 | 0.83 | 0.12 |
| | Control& | P value | <0.0001 | <0.0001 | 0.14 | 0.03 | 0.21 |
| | | Summary | **** | **** | ns | **** | ns |
| Day12 | Mean | 0.45 | 0.58 | 5.35 | 0.25 | 0.35 | 0.05 |
| | SD | 0.83 | 0.53 | 0.40 | 0.12 | 0.59 | 0.05 |
| | Control& | P value | 0.028 | <0.0001 | 0.37 | 0.66 | 0.04 |
| | | Summary | * | **** | ns | ns | * |

**Table S2.** Multicellular Mass Counts in Drug-Treated Groups Versus Controls (Day 12, n=20).

| | Control | VCR | PTX | DDP | DOX | 5-FU |
|---|---|---|---|---|---|---|
| Mean | 0.27 | 8.50 | 36.17 | 1.83 | 0.50 | 0.35 |
| SD | 0.15 | 0.76 | 1.80 | 0.48 | 0.43 | 0.16 |
| Control&drug | P value | <0.0001 | <0.0001 | 0.018 | 0.022 | 0.72 |

| | | | | | | |
|---|---|---|---|---|---|---|
| | Summary | **** | **** | * | * | ns |
| PTX& | P value | <0.0001 | | <0.0001 | <0.0001 | <0.0001 |
| | Summary | **** | | **** | **** | **** |

**Table S3.** Giant multicellular mass formation: 48-h pulse (10 μM DOX/PTX) + 10-day recovery versus continuous 12-day exposure (n=6).

| | | Sustained drug | Withdraw drug | P value | Summary |
|---|---|---|---|---|---|
| VINC | Mean | 0.55 | 4.55 | <0.0001 | **** |
| | SEM | 0.14 | 0.35 | | |
| PTX | Mean | 5.35 | 6.05 | 0.22 | ns |
| | SEM | 0.40 | 0.40 | | |

**Table S4.** Cell viability following treatment with the indicated drugs (n=8).

| | VCR | | PTX | | DDP | | DOX | | 5-FU | |
|---|---|---|---|---|---|---|---|---|---|---|
| | Mean | SD | Mean | SD | Mean | SD | Mean | SD | Mean | SD |
| Day0 | 1 | 0.093 | 1 | 0.093 | 1 | 0.093 | 1 | 0.093 | 1 | 0.093 |
| Day1 | 1.07 | 0.078 | 1.09 | 0.16 | 1.19 | 0.10 | 0.98 | 0.11 | 1.04 | 0.11 |
| Day2 | 0.76 | 0.14 | 0.98 | 0.099 | 0.99 | 0.064 | 0.80 | 0.14 | 0.82 | 0.14 |
| Day4 | 0.64 | 0.079 | 0.69 | 0.082 | 0.68 | 0.073 | 0.22 | 0.034 | 0.47 | 0.070 |

| | | | | | | | | | |
|---|---|---|---|---|---|---|---|---|---|
| Day8 | 0.097 | 0.028 | 0.32 | 0.097 | 0.080 | 0.023 | 0.052 | 0.021 | 0.17 | 0.022 |
| Day12 | 0.041 | 0.019 | 0.17 | 0.039 | 0.016 | 0.013 | 0.011 | 0.0081 | 0.10 | 0.027 |

**Table S5.** The survival ratios (%) of cells treated with different drugs for 12 days (n=3).

| | VCR | PTX | DDP | DOX | 5-FU |
|---|---|---|---|---|---|
| Multicellular cells | 96 | 98 | 99 | 80.7 | 61 |
| Big mononuclear cells | 2.8 | 1.9 | 0.95 | 29.3 | 30.5 |
| Small cells | 1.2 | 0.1 | 0.05 | 3 | 8.5 |

**Table S6.** Unicell-to-daughter cell ratio following spheroid replating (6-day PTX pretreatment, days 3 and 6, n=3).

| | PTX 1 μM | | PTX 5μM | | PTX 10 | | PTX 1 μM& 5μM | | PTX 1 μM& 10μM | | PTX 5 μM& 10μM | |
|---|---|---|---|---|---|---|---|---|---|---|---|---|
| | Mean | SEM | Mean | SEM | Mean | SEM | P value | Summary | P value | Summary | P value | Summary |
| Day3 | 83.67 | 2.33 | 11.00 | 1.53 | 6.00 | 6.58 | <0.0001 | **** | <0.0001 | **** | 0.037 | * |
| Day6 | 93 | 1.52 | 24.67 | 1.30 | 14.33 | 1.20 | <0.0001 | **** | <0.0001 | **** | 0.0037 | *** |

**Table S7.** Daughter cell generation capacity post-PTX withdrawal (0–10 μM, Various Durations) (n=3).

| | treatment | Day5 | Day 16 | Day 21 |
|---|---|---|---|---|

| Time of withdraw | | 1 μM | 5 μM | 10 μM | 1 μM | 5 μM | 10 μM | 1 μM | 5 μM | 10 μM | Day5& Day 16 | Day5& Day 21 | Day16& Day 21 |
|---|---|---|---|---|---|---|---|---|---|---|---|---|---|
| D15 | Mean | 12.80 | 4.23 | 1.83 | 2.37 | 0.00 | 0.00 | 0.00 | 0.00 | 0.00 | 1 μM: 0.002, ***<br>5 μM: 0.005, **<br>10 μM: 0.02, * | 1 μM: 0.0009, ***<br>5 μM: 0.005, **<br>10 μM: 0.02, * | 1 μM: 0.01, **<br>5 μM: ns<br>10 μM: ns |
| | SEM | 1.45 | 0.79 | 0.49 | 0.52 | 0.00 | 0.00 | 0.00 | 0.00 | 0.00 | | | |
| | P value, Summary | 1 μM&5 μM: 0.006, **<br>1 μM&10 μM: 0.002, **<br>5 μM&10 μM: 0.06, ns | | | 1 μM&5 μM: 0.010455<br>1 μM&10 μM: 0.010455<br>5 μM&10 μM: ns | | | 1 μM&5 μM: ns<br>1 μM&10 μM: ns<br>5 μM&10 μM: ns | | | | | |
| D21 | Mean | 36.43 | 31.20 | 36.00 | 21.90 | 4.50 | 1.03 | 0.20 | 0.00 | 0.00 | 1 μM: 0.007, ***<br>5 μM: <0.0001, ****<br>10 μM: <0.0001, **** | 1 μM: <0.0001, ****<br>5 μM: <0.0001, ****<br>10 μM: <0.0001, **** | 1 μM: 0.0003, ***<br>5 μM: 0.002, **<br>10 μM: 0.02, * |
| | SEM | 2.17 | 1.57 | 2.08 | 1.86 | 0.62 | 0.29 | 0.15 | 0.00 | 0.00 | | | |
| | P value, Summary | 1 μM&5 μM: 0.12, ns<br>1 μM&10 μM: 0.89, ns | | | 1 μM&5 μM: 0.0008, ***<br>1 μM&10 μM: 0.0003, | | | 1 μM&5 μM: 0.26, ns<br>1 μM&10 μM: 0.26, ns | | | | | |

| | | | | | | | | | | | | | |
|---|---|---|---|---|---|---|---|---|---|---|---|---|---|
| | | 5 µM&10 µM: 0.14, ns | | | ***<br>5 µM&10 µM: 0.007, ** | | | 5 µM&10 µM: ns | | | | | |
| D35 | Mean | 66.00 | 62.07 | 60.77 | 53.27 | 50.67 | 32.60 | 17.00 | 10.53 | 9.57 | 1 µM: 0.03, *<br>5 µM: 0.019, *<br>10 µM: 0.000488 | 1 µM: <0.0001, ****<br>5 µM: <0.0001, ****<br>10 µM: <0.0001, **** | 1 µM: <0.0001<br>5 µM: 0.00013<br>10 µM: 0.000980 |
| | SEM | 2.01 | 1.94 | 1.60 | 3.40 | 2 33 | 2.19 | 0.79 | 1.49 | 1.51 | | | |
| | P value, Summary | 1 µM&5 µM: 0.23, ns<br>1 µM&10 µM: 0.11, ns<br>5 µM&10 µM: 0.63, ns | | | 1 µM&5 µM: 0.56, ns<br>1 µM&10 µM: 0.007, **<br>5 µM&10 µM: 0.005, ** | | | 1 µM&5 µM: 0.018, *<br>1 µM&10 µM: 0.012, *<br>5 µM&10 µM: 0.67, ns | | | | | |
| D45 | Mean | | | | 62.07 | 62.87 | 59.37 | 59.07 | 57.33 | 36.97 | | | 1 µM: 0.27, ns<br>5 µM: 0.14, ns<br>10 µM: |
| | SEM | | | | 1.25 | 1.43 | 3.55 | 2.00 | 2.71 | 3.17 | | | |

| | | | | | | | | | | | | | |
|---|---|---|---|---|---|---|---|---|---|---|---|---|---|
| | | | | | | | | | | | | | 0.009, ** |
| D15&D21 | P value, Summary | | | | 1 µM&5 µM: 0.69, ns<br>1 µM&10 µM: 0.51, ns<br>5 µM&10 µM: 0.41, ns | | | 1 µM&5 µM: 0.63, ns<br>1 µM&10 µM: 0.004, ns<br>5 µM&10 µM: 0.008, ns | | | | | |

**Table S8.** Daughter cells derived from PTX-treated 4T1 cells.

| Cell name | Suspending/attached | Duration of PTX exposure (Days) |
|---|---|---|
| TS1 | Suspending spheres | 6 |
| TS2 | Suspending spheres | 6 |
| TS3 | Suspending spheres | 6 |
| TS4 | Suspending spheres | 3 |
| TS5 | Suspending spheres | 3 |
| TS6 | Suspending spheres | 3 |
| TA1 | Attached cells | 21 |
| TA2 | Attached cells | 21 |

| | | |
|---|---|---|
| TA3 | Attached cells | 21 |
| TA4 | Attached cells | 16 |
| TA5 | Attached cells | 16 |
| TA6 | Attached cells | 16 |
| TA7 | Attached cells | 5 |
| TA8 | Attached cells | 5 |
| TA9 | Attached cells | 5 |

**Table S9.** Growth capacity in control and PTX-pretreated cultures (n=5).

| | Control | | PTX-day2 | | PTX-day14 | | |
|---|---|---|---|---|---|---|---|
| | Mean | SEM | Mean | SEM | Mean | SEM | PTX-day 2 & Control |
| 8 | 59.77 | 5.079 | 0 | 0 | 0 | 0 | p<0.0001 |
| 10 | 72.78 | 13.904 | 13.238 | 2.529 | 0 | 0 | **** |
| 12. | 138.06 | 45.526 | 23.798 | 4.306 | 0 | 0 | |
| 14 | 232.82 | 73.854 | 40.578 | 9.505 | 0 | 0 | PTX-day 14 & Control |
| 16 | 270.61 | 76.653 | 74.490 | 10.534 | 0 | 0 | p<0.0001 |

| | | | | | | | |
|---|---|---|---|---|---|---|---|
| 18 | 409.06 | 112.812 | 88.214 | 13.126 | 0 | 0 | **** |
| 20 | 551.12 | 132.090 | 119.160 | 22.396 | 0 | 0 | |
| 22 | 663.61 | 205.498 | 153.902 | 28.470 | 0 | 0 | PTX-day 2 & PTX-day |
| 24 | 778.29 | 190.855 | 186.610 | 27.279 | 0 | 0 | p<0.0001 |
| 26 | 924.92 | 225.427 | 268.272 | 33.885 | 0 | 0 | **** |
| 28 | 983.88 | 244.720 | 388.520 | 68.715 | 0 | 0 | |
| 30 | 1100.80 | 275.359 | 453.060 | 87.110 | 0 | 0 | |
| 37 | | | | | 10.72 | 10.724 | |
| 41 | | | | | 150.77 | 98.766 | |
| 45 | | | | | 277.06 | 261.545 | |
| 48 | | | | | 565.99 | 369.216 | |
| 51 | | | | | 965.08 | 684.264 | |

**Table S10.** Large round cell formation capacity upon PTX treatment (2 days) in daughter cells derived from drug-withdrawn cultures compared to standard 4T1 cultures (n=20).

| | | Control | PTX-10 μM | Control& PTX |
|---|---|---|---|---|

| | | | | |
|---|---|---|---|---|
| 4T1 | Mean | 1.00 | 98.67 | <0.0001**** |
| | SEM | 0.25 | 1.72 | |
| TS6 | Mean | 12.89 | 100.78 | <0.0001**** |
| | SEM | 0.77 | 1.09 | |
| | Control& | <0.0001**** | ns | |
| TA3 | Mean | 10.33 | 99.44 | <0.0001**** |
| | SEM | 0.49 | 1.16 | |
| | Control& | <0.0001**** | ns | |

**Table S11.** $AP^+$ Cell formation capacity: control versus PTX-treated groups (n=30).

| | | Control | 1 µM | 5 µM | 10 µM | 1 µM&5 µM | 1 µM & 10 µM | 1 µM&10 µM |
|---|---|---|---|---|---|---|---|---|
| 0 | Mean | 21.17 | 21.33 | 21.00 | 21.33 | | | |
| | SEM | 0.48 | 0.33 | 0.52 | 0.422 | | | |
| 4 | Mean | 15.17 | 20.00 | 25.33 | 30.00 | <0.0001 **** | <0.0001**** | 0.00017*** |
| | SEM | 0.601 | 0.52 | 0.49 | 0.63 | | | |
| | Drug | | 0.00012*** | <0.0001**** | <0.0001**** | | | |

| | | | | | | | | |
|---|---|---|---|---|---|---|---|---|
| | &Control | | | | | | | |
| 8 | Mean | 19 | 29.17 | 58.33 | 60.83 | <0.0001**** | <0.0001**** | 0.0045** |
| | SEM | 0.58 | 0.60 | 0.49 | 0.48 | | | |
| | Drug &Control | | <0.0001**** | <0.0001**** | <0.0001**** | | | |
| 12 | Mean | 25 | 42.17 | 59.50 | 56.00 | <0.0001**** | <0.0001**** | 0.001** |
| | SEM | 0.516 | 0.60 | 0.43 | 0.63 | | | |
| | Drug &Control | | <0.0001**** | <0.0001**** | <0.0001**** | | | |
| 24 | Mean | 62 | 75.17 | 75.50 | 58.17 | 0.66, ns | <0.0001**** | <0.0001**** |
| | SEM | 0.58 | 0.42 | 0.56 | 0.48 | | | |
| | Drug &Control | | <0.0001**** | <0.0001**** | 0.00452** | | | |

**Table S12.** Quantification of AP-positive cell ratios at different time points comparing PTX-treated and untreated groups (n=30).

| | | Control | 1 μM | 5 μM | 10 μM | 1 μM&5 μM | 1 μM & 10 μM | 5 μM&10 μM |
|---|---|---|---|---|---|---|---|---|
| 0 | Mean | 16.67 | 16.50 | 16.50 | 16.33 | | | |

| | | | | | | | | |
|---|---|---|---|---|---|---|---|---|
| | SEM | 0.76 | 0.76 | 0.62 | 0.88 | | | |
| 4 | Mean | 9.48 | 15.55 | 20.02 | 23.17 | <0.0001 **** | <0.0001**** | <0.0001**** |
| | SEM | 0.41 | 0.33 | 0.29 | 0.26 | | | |
| | Drug &Control | | <0.0001**** | <0.0001**** | <0.0001**** | | | |
| 8 | Mean | 9.72 | 16.20 | 33.37 | 44.03 | <0.0001**** | <0.0001**** | <0.0001**** |
| | SEM | 0.22 | 0.26 | 0.20 | 0.41 | | | |
| | Drug &Control | | <0.0001**** | <0.0001**** | <0.0001**** | | | |
| 12 | Mean | 12.40 | 23.72 | 39.58 | 41.20 | <0.0001**** | <0.0001**** | 0.0038** |
| | SEM | 0.30 | 0.26 | 0.32 | 0.29 | | | |
| | Drug &Control | | <0.0001**** | <0.0001**** | <0.0001**** | | | |
| 24 | Mean | 20.01 | 32.25 | 42.13 | 40.82 | <0.0001**** | <0.0001**** | 0.0047** |
| | SEM | 0.28 | 0.21 | 0.24 | 0.27 | | | |
| | Drug &Control | | <0.0001**** | <0.0001**** | <0.0001**** | | | |

**Table S13**. Quantification of AP-positive large round cell formation at different time points comparing PTX-treated and untreated groups (n=30).

| | | Control | 1 μM | 5 μM | 10 μM | 1 μM&5 μM | 1 μM & 10 μM | 5 μM&10 μM |
|---|---|---|---|---|---|---|---|---|
| 0 | Mean | 1.000 | 1.000 | 1.000 | 1.000 | | | |
| | SEM | 0.028 | 0.013 | 0.015 | 0.009 | | | |
| 4 | Mean | 1.234 | 1.110 | 1.110 | 1.032 | <0.0001 **** | <0.0001**** | 0.00012 *** |
| | SEM | 0.015 | 0.015 | 0.015 | 0.014 | | | |
| | Drug &Control | | | | | | | |
| 8 | Mean | 1.411 | 1.275 | 1.149 | 1.100 | <0.0001**** | <0.0001**** | 0.025* |
| | SEM | 0.015 | 0.020 | 0.015 | 0.014 | | | |
| | Drug &Control | | | | | | | |
| 12 | Mean | 1.649 | 1.409 | 1.251 | 1.118 | <0.0001**** | <0.0001**** | <0.0001**** |
| | SEM | 0.016 | 0.017 | 0.016 | 0.017 | | | |
| | Drug &Control | | | | | | | |
| 24 | Mean | 2.275 | 1.576 | 1.251 | 1.102 | <0.0001**** | <0.0001**** | <0.0001**** |
| | SEM | 0.029 | 0.023 | 0.019 | 0.018 | | | |
| | Drug &Control | | | | | | | |

**Table S14.** Quantification of AP-positive oocyte-like cell formation at different time points comparing PTX-treated and untreated groups (n=30).

| | | Control | 1 μM | 5 μM | 10 μM | 1 μM&5 μM | 1 μM & 10 μM | 1 μM&10 μM |
|---|---|---|---|---|---|---|---|---|
| 0 | Mean | 1 | 1 | 1 | 1 | | | |
| | SEM | 0.246 | 0.211 | 0.178 | 0.205 | | | |
| 4 | Mean | 1.071 | 23.714 | 29.286 | 31.714 | <0.0001 **** | <0.0001**** | 0.0099 ** |
| | SEM | 0.199 | 0.901 | 0.628 | 0.661 | | | |
| | Drug &Control | | <0.0001**** | <0.0001**** | <0.0001**** | | | |
| 8 | Mean | 1.429 | 40.417 | 74.500 | 94.929 | <0.0001**** | <0.0001**** | <0.0001**** |
| | SEM | 0.188 | 1 291 | 1.667 | 4.342 | | | |
| | Drug &Control | | <0.0001**** | <0.0001**** | <0.0001**** | | | |
| 12 | Mean | 0.929 | 40.571 | 81.500 | 74.429 | <0.0001**** | <0.0001**** | 0.0034 ** |
| | SEM | 0.197 | 1.471 | 1.972 | 1.220 | | | |
| | Drug &Control | | <0.0001**** | <0.0001**** | <0.0001**** | | | |

| | | | | | | | | |
|---|---|---|---|---|---|---|---|---|
| 24 | Mean | 0.857 | 45.071 | 51.143 | 44.571 | 0.027 * | 0.77 ns | 0.0098** |
| | SEM | 0.195 | 1.418 | 2.269 | 0.955 | | | |
| | Drug &Control | | <0.0001**** | <0.0001**** | <0.0001**** | | | |

**Table S15.** Relative mRNA expression levels of meiosis-related genes in 4T1 cells treated with 10 μM PTX versus untreated controls (n=3).

| | | 0h | 4h | | 8h | | 12h | | 24h | |
|---|---|---|---|---|---|---|---|---|---|---|
| | | | Control | PTX | Control | PTX | Control | PTX | Control | PTX |
| Blimp1 | Mean | 1.020 | 1.569 | 1.831 | 6.638 | 6.206 | 16.479 | 5.707 | 2.972 | 3.617 |
| | SEM | 0.137 | 0.307 | 0.44 | 1.999 | 0.950 | 4.575 | 0.973 | 0.771 | 1.121 |
| | p value | | 0.650 | | 0.855 | | 0.083 | | 0.653 | |
| | Summary | | ns | | ns | | ns | | ns | |
| IFITM3 | Mean | 1.004 | 3.533 | 3.119 | 4.029 | 3.147 | 3.123 | 1.175 | 1.605 | 0.974 |
| | SEM | 0.059 | 0.065 | 0.385 | 0.018 | 0.056 | 0.116 | 0.215 | 0.355 | 0.028 |
| | p value | | 0.3477 | | 0.0001 | | 0.0013 | | 0.151 | |
| | Summary | | ns | | *** | | ** | | ns | |

| | | | | | | | | | | |
|---|---|---|---|---|---|---|---|---|---|---|
| Oct4 | Mean | 1.045 | 10.272 | 5.476 | 7.132 | 45.62 | 1.532 | 5.561 | 5.176 | 23.78 |
| | SEM | 0.200 | 0.512 | 0.83 | 1.04 | 1.92 | 5.56 | 0.348 | 0.119 | 2.358 |
| | p value | | 0.008 | | <0.0001 | | 0.0008 | | 0.0014 | |
| | Summary | | ** | | **** | | *** | | ** | |
| Sox2 | Mean | 1.090 | 16.74 | 6.23 | 10.94 | 70.86 | 7.861 | 3.914 | 3.543 | 10.78 |
| | SEM | 0.278 | 4.98 | 1.75 | 2.60 | 7.86 | 2.433 | 1.54 | 0.598 | 1.08 |
| | p value | | 0.117 | | <0.0001 | | 0.242 | | 0.0043 | |
| | Summary | | ns | | **** | | ns | | ** | |
| Stellar | Mean | 1.018 | 1.98 | 1.48 | 2.348 | 10.89 | 0.619 | 1.086 | 1.324 | 6.95 |
| | SEM | 0.13 | 0.081 | 0.016 | 0.085 | 0.392 | 0.057 | 0.459 | 0.129 | 0.397 |
| | p value | | 0.0039 | | <0.0001 | | 0.397 | | 0.0002 | |
| | Summary | | ** | | **** | | ns | | *** | |
| MYC | Mean | 1.003 | 2.168 | 1.479 | 1.975 | 1.944 | 1.950 | 0.521 | 0.680 | 0.407 |
| | SEM | 0.053 | 0.100 | 0.196 | 0.357 | 0.035 | 0.307 | 0.072 | 0.020 | 0.079 |
| | p value | | 0.0352 | | 0.935 | | 0.011 | | 0.029 | |
| | Summary | | * | | ns | | * | | * | |

| | | | | | | | | | | |
|---|---|---|---|---|---|---|---|---|---|---|
| STAT3 | Mean | 1.007 | 2.455 | 2.267 | 2.178 | 1.667 | 1.47 | 0.768 | 1.571 | 1.431 |
| | SEM | 0.087 | 0.020 | 0.071 | 0.054 | 0.131 | 0.231 | 0.273 | 0.049 | 0.099 |
| | p value | | 0.064 | | 0.022 | | 0.120 | | 0.273 | |
| | Summary | | ns | | * | | ns | | ns | |
| Nanog | Mean | 1.005 | 5.63 | 4.08 | 4.09 | 24.49 | 1.11 | 3.50 | 3.31 | 16.29 |
| | SEM | 0.07 | 0.44 | 0.73 | 0.23 | 3.04 | 0.031 | 0.42 | 0.43 | 0.72 |
| | p value | | 0.144 | | 0.0026 | | 0.0062 | | <0.0001 | |
| | Summary | | ns | | ** | | ** | | **** | |
| NANOS3 | Mean | 1.006 | 1.025 | 1.465 | 1.046 | 0.778 | 0.665 | 0.652 | 0.336 | 0.569 |
| | SEM | 0.08 | 0.048 | 0.072 | 0.067 | 0.065 | 0.034 | 0.033 | 0.014 | 0.044 |
| | p value | | 0.0069 | | 0.0456 | | 0.8639 | | 0.0074 | |
| | Summary | | *** | | * | | ns | | ** | |

**Table S16.** Estrogen concentration dynamics in control versus drug-treated groups.

| | Control | PTX |
|---|---|---|
| 0 | 101 | 101 |
| Day1 | 1164 | 985 |
| Day3 | 3229 | 3363 |
| Day4 | 4774 | 5338 |
| Day6 | | 5455 |

**Table S17.** Relative mRNA expression levels of meiosis-related genes in 4T1 cells treated with 10 μM PTX compared to untreated control group (n=3).

| | | 0h | Day 1 | | Day 2 | | Day 3 | | Day 4 | | |
|---|---|---|---|---|---|---|---|---|---|---|---|
| | | | Control | PTX | Control | PTX | Control | PTX | Control | PTX | PTX-upper |
| DMC1 | Mean | 1.08 | 1.13 | 13.15 | 5.70 | 11.76 | 4.11 | 18.12 | 2.85 | 26.79 | 260.63 |
| | SEM | 0.48 | 0.27 | 5.66 | 1.82 | 8.61 | 3.19 | 3.46 | 1.23 | 4.85 | 98.29 |
| | p value | | 0.021 | | 0.299 | | 0.0067 | | 0.0012 | | PTX & PTX-upper |
| | | | | | | | | | | | 0.015 |
| | Summary | | * | | ns | | ** | | ** | | * |
| SYCP3 | Mean | 1.00 | 0.72 | 2.28 | 0.56 | 6.23 | 0.57 | 19.10 | 1.79 | 13.99 | 127.85 |

| | | | | | | | | | | | |
|---|---|---|---|---|---|---|---|---|---|---|---|
| | SEM | 0.084 | 0.37 | 1.81 | 0.40 | 1.32 | 0.46 | 10.66 | 0.37 | 4.23 | 52.31 |
| | p value | | 0.217 | | 0.0021 | | 0.039 | | 0.0076 | | PTX & PTX-upper<br>0.020 |
| | Summary | | | | | | | | | | * |

**Table S18.** Dose- and time-dependent quantification of alkaline phosphatase (AP)-positive blastomere-like cells in 4T1 cultures treated with PTX compared to control groups (n=9).

| | | Control | 0.5 µM | 1 µM | 2 µM | 5 µM | 10 µM | 0.5µM& others | 1 µM & others | others |
|---|---|---|---|---|---|---|---|---|---|---|
| 0 | Mean | 1.000 | 1.000 | 1.000 | 1.000 | 1.000 | 1.000 | | | |
| | SEM | 0.013 | 0.015 | 0.016 | 0.014 | 0.009 | 0.013 | | | |
| 1 | | 1.185 | 22.000 | 40.550 | 81.050 | 102.750 | 155.717 | & 1 µM <0.0001****<br>& 2 µM <0.0001****<br>& 5 µM<0.0001****<br>& 10 µM <0.0001**** | & 2 µM <0.0001****<br>& 5 µM<0.0001****<br>& 10 µM<0.0001**** | 2 µM & & 5 µM <0.0001****<br>2 µM & & 10 µM <0.0001****<br>5 µM & & 10 µM <0.0001**** |
| | | 0.018 | 0.419 | 0.407 | 0.300 | 0.519 | 0.459 | | | |
| | | <0.0001 | <0.0001 | <0.0001 | <0.0001 | <0.0001 | <0.0001 | | | |
| | | **** | **** | **** | **** | **** | **** | | | |

| | | | | | | | | | | |
|---|---|---|---|---|---|---|---|---|---|---|
| 2 | Mean | 1.243 | 36.067 | 59.083 | 72.183 | 101.517 | 154.633 | & 1 μM <0.0001****<br>& 2 μM <0.0001****<br>& 5 μM<0.0001****<br>& 10 μM <0.0001**** | & 2 μM <0.0001****<br>& 5 μM<0.0001****<br>& 10 μM<0.0001**** | 2 μM & & 5 μM <0.0001****<br>2 μM & & 10 μM <0.0001****<br>5 μM & & 10 μM <0.0001**** |
| | SEM | 0.024 | 0.346 | 0.315 | 0.318 | 0.285 | 0.448 | | | |
| | Drug &Control | <0.0001 | <0.0001 | <0.0001 | <0.0001 | <0.0001 | <0.0001 | | | |
| | | **** | **** | **** | **** | **** | **** | | | |
| 3 | Mean | 1.255 | 36.850 | 65.467 | 83.967 | 100.250 | 80.700 | & 1 μM <0.0001****<br>& 2 μM <0.0001****<br>& 5 μM<0.0001****<br>& 10 μM <0.0001**** | & 2 μM <0.0001****<br>& 5 μM<0.0001****<br>& 10 μM<0.0001**** | 2 μM & & 5 μM <0.0001****<br>2 μM & & 10 μM <0.0001****<br>5 μM & & 10 μM <0.0001**** |
| | SEM | 0.030 | 0.453 | 0.357 | 0.400 | 0.467 | 0.268 | | | |
| | Drug &Control | <0.0001 | <0.0001 | <0.0001 | <0.0001 | <0.0001 | <0.0001 | | | |
| | | **** | **** | **** | **** | **** | **** | | | |
| 4 | Mean | 1.172 | 50.367 | 43.617 | 46.367 | 56.383 | 55.133 | & 1 μM <0.0001****<br>& 2 μM <0.0001****<br>& 5 μM<0.0001****<br>& 10 μM | & 2 μM 0.0004***<br>& 5 μM<0.0001****<br>& 10 μM<0.0001**** | 2 μM & & 5 μM <0.0001****<br>2 μM & & 10 μM |
| | SEM | 0.043 | 0.282 | 0.374 | 0.378 | 0.275 | 0.409 | | | |
| | Drug &Control | <0.0001 | <0.0001 | <0.0001 | <0.0001 | <0.0001 | <0.0001 | | | |

| | | | | | | | | | | |
|---|---|---|---|---|---|---|---|---|---|---|
| | | **** | **** | **** | **** | **** | **** | <0.0001**** | | <0.0001****<br>5 µM & & 10 µM<br>0.029 * |
| 5 | Mean | 1.127 | 9.033 | 26.567 | 26.567 | 21.050 | 26.317 | & 1 µM <0.0001****<br>& 2 µM <0.0001****<br>& 5 µM<0.0001****<br>& 10 µM <0.0001**** | & 2 µM >0.99 ns<br>& 5 µM 0.257 ns<br>& 10 µM 0.567 ns | 2 µM & & 5 µM<br>0.258 ns<br>2 µM & & 10 µM<br>0.057 ns<br>5 µM & & 10 µM<br>0.48 ns |
| | SEM | 0.038 | 0.364 | 0.334 | 0.334 | 0.315 | 0.257 | | | |
| | Drug &Control | <0.0001 | <0.0001 | <0.0001 | <0.0001 | <0.0001 | <0.0001 | | | |
| | | **** | **** | **** | **** | **** | **** | | | |

**Table S19.** Comparative capacity for large round cell formation induced by 10 µM PTX between 4T1 and 168FARN cultures (n=3).

| | 4T1 | | 168FARN | | 4T1 & 168FARN | |
|---|---|---|---|---|---|---|
| | Mean | SEM | Mean | SEM | P value | Summary |
| 0 | 1.00 | 0.350 | 0.33 | 0.229 | 0.119 | ns |
| 48h | 91.67 | 2.030 | 1.17 | 0.365 | <0.0001 | **** |
| 192h | 30.67 | 1.270 | 0.17 | 0.167 | <0.0001 | **** |

**Table S20.** Differential capacity for large round cell formation induced by 10 μM PTX between control 4T1 cells and ACVR1 or DAZL knockout derivatives (n=20).

| | Guide | | KO-ACVR1 | | KO-DAZL | | Guide & KO-ACVR1 | | Guide & KO- DAZL | |
|---|---|---|---|---|---|---|---|---|---|---|
| | Mean | SEM | Mean | SEM | Mean | SEM | P value | Summary | P value | Summary |
| 48h | 1.000 | 0.014 | 0.018 | 0.006 | 0.020 | 0.005 | <0.0001 | **** | <0.0001 | **** |
| 192h | 0.465 | 0.015 | 0.002 | 0.002 | 0.007 | 0.004 | <0.0001 | **** | <0.0001 | **** |

**Table S21.** Capacity for large round cell formation in 4T1 cultures following various drug treatments (n=6).

| | | Control | PTX | LDN212854 | PTX+LDN212854 | LDN193189 | PTX+LDN193189 |
|---|---|---|---|---|---|---|---|
| 0 | Mean | 1.000 | 1.000 | 1.000 | 1.000 | 1.000 | 1.000 |
| | SEM | 0.350 | 0.29 | 0.27 | 0.33 | 0.37 | 0.28 |
| 24 | Mean | 1.333 | 98.000 | 0.667 | 0.833 | 0.682 | 1.333 |
| | SEM | 0.507 | 2.719 | 0.306 | 0.410 | 0.305 | 0.446 |
| 48h | Mean | 1.579 | 91.667 | 0.333 | 0.167 | 0.167 | 0.00 |
| | SEM | 0.468 | 2.030 | 0.229 | 0.167 | 0.167 | 0.00 |
| PTX & LDN212854 p<0.0001****, PTX & PTX+LDN212854 p<0.0001****, PTX & LDN193189 p<0.0001****, PTX & | | | | | | | |

| |
|---|
| PTX+LDN193189 p<0.0001****, LDN212854&PTX+LDN212854 p=0.148ns, LDN193189 & PTX+ LDN193189 p=0.012* |

**Table S22.** Survival of large round cells derived from 5-day PTX-treated 4T1 cultures following drug withdrawal and exposure to indicated compounds (n=20).

| | Control | | LDN212854 | | LDN193189 | | Control & LDN212854 | | Control & LDN193189 | |
|---|---|---|---|---|---|---|---|---|---|---|
| | Mean | SEM | Mean | SEM | Mean | SEM | P value | Summary | P value | Summary |
| 0 | 1.000 | 0.045 | 1.000 | 0.043 | 1.000 | 0.038 | | ns | | ns |
| 48h | 0.891 | 0.035 | 0.711 | 0.046 | 0.011 | 0.006 | 0.034 | * | <0.0001 | **** |
| 72h | 0.842 | 0.043 | 0.035 | 0.011 | 0.007 | 0.005 | <0.0001 | **** | <0.0001 | **** |
| 144h | 0.810 | 0.036 | 0.007 | 0.005 | 0.004 | 0.004 | <0.0001 | **** | <0.0001 | **** |

**Table S23.** Primers for qRT-PCR and nested PCR.

| **Gene** | **Forward/reverse** | **Sequence 5'-3'** | **Size (bp)** |
|---|---|---|---|
| ***OCT4*** | forward | TCAGGTTGGACTGGGCCTAGT | 100 |
| | reverse | GGAGGTTCCCTCTGAGTTGCTT | |
| ***hqSOX2*** | forward | GAGGGCTGGACTGCGAACT | 76 |
| | reverse | TTTGCACCCCTCCCAATTC | |
| ***hqNANOG*** | forward | GAAATCCCTTCCCTCGCCATC | 160 |
| | reverse | CTCAGTAGCAGACCCTTGTAAGC | |

| | | | |
|---|---|---|---|
| ***mqIFITM3*** | forward | GTTATCACCATTGTTAGTGTCATC | 151 |
| | reverse | AATGAGTGTTACACCTGCGTG | |
| ***mqSTELLAR*** | forward | GCAGTCTACGGAACCGCATT | 123 |
| | reverse | GGTCTTTCAGCACCGACAACA | |
| ***mqNANOS3*** | forward | TAAGGCTGGATCCCAAACCA | 115 |
| | reverse | GACTCGCCATTGTGTTTGCA | |
| ***mqSYCP3*** | forward | GCAATCAAACAGATACACGAG | 144 |
| | reverse | CTGCTGAGTTTCCATCATAAC | |
| ***mqBLIMP1*** | forward | AGCATGACCTGACATTGACACC | 162 |
| | reverse | CTCAACACTCTCATGTAAGAGGC | |
| ***mqKLF4*** | forward | GCACACCTGCGAACTCACAC | 53 |
| | reverse | CCGTCCCAGTCACAGTGGTAA | |
| ***mqDMC1*** | forward | GATCCAGGAGCAACTATGACC | 162 |
| | reverse | GGCTTCATTTTCAGGCATCTCG | |
| ***nqZP3*** | forward | GATCCCCGATAAGCTCAACA | 283 |
| | reverse | GGTTTGAGCAGAAGCAGTCC | |
| ***mqGADPH*** | forward | AAGGGCTCATGACCACAGTC | 207 |
| | reverse | ACACATTGGGGGTAGGAACA | |
| ***mqMYC*** | forward | TCTCCACTCACCAGCACAACTACG | 103 |
| | reverse | ATCTGCTTCAGGACCCT | |
| ***mH19DMR*** | outside forward | GAGTATTTAGGAGGTATAAGAATT | |
| | outside reverse | **ATCAAAAACTAACATAAACCCCT** | |
| | inside forward | GTAAGGAGATTATGTTTATTTTTGG | |

| | | |
|---|---|---|
| | inside reverse | CCTCATTAATCCCATAACTAT |

**Table S24.** sgRNAs sequences (Sigma).

| Ensemble Symbol | Clone ID | chr_start | chr_end seq | target with PAM |
|---|---|---|---|---|
| Dazl | MM5000020903 | 50289105 | 50289126 | CAAACACCGTTTTTGTTGGAGG |
| Acvr1 | MM5000002095 | 58479837 | 58479858 | ACACTGCTGGCCTTCACAGTGG |

**Table S25.** The primers were used for the Sanger Sequencing of CRSPR-Cas9 knockout.

| Gene | Sequence 5'-3' |
|---|---|
| ACVR1 | TGTCAGGAGTCTGGGCATCA |
| DAZL | TGCTTTCAGTTTCAGAGGTAA |

**Table S26.** Sequencing data of knockout cells generated by CRISPR-Cas9 technology.

| | |
|---|---|
| 4T1-KO-Acvr1-1# | Query 241 AGGTCAACCAGAAACTTTACAtg----tgtgtg-------------------------- 269<br>\|\|\|\|\|\|\|\|\|\|\|\|\|\|\|\|\|\|\|\|\| \|\|\|\|\|\| |

<table>
<tr><td></td><td>Sbjct 58479916 AGGTCAACCAGAAACTTTACATGTGTGTGTGAGGGCCTCTCCTGCGGGAACGAGGACC 58479857<br><br>Query 270 --tgtgAGGGCC-----------TCTCCTGCGGG----AACGAGGAC---CAC-TGT--- 305<br>||||| |||| ||| || | | ||||| | | ||| | |<br>Sbjct 58479856 ACTGTGAAGGCCAGCAGTGTTTTTCTTCTCTGAGCATCAACGATGGCTTCCACGTCTACC 58479797<br>sgRNA ACACTGCTGGCCTTCACAGTGG Del:58<br>Query 306 ----GGGCTGCTTTCAGGTTTATTAGCAGGGGAAGATGACGTGTAACACCCCGCCGTCAC 361<br>||||||||||||||||||| |||||||||||||||||||||| |||||||||||||<br>Sbjct 58479796 AGAAGGGCTGCTTTCAGGTTTATGAGCAGGGGAAGATGACGTGTAAGACCCCGCCGTCAC 58479737<br>sgRNA AACCTGAAAGCAGCCCTTCTGG</td></tr>
<tr><td>4T1-KO-Acvr1-2#</td><td>Query 241 AGGTCAACCAGAAACTTTACAtgtgtgtgtgtgAGGGCCTCTCCTGCGGGAACG------ 294<br>||||||||||||||||||||||||||||||||||||||||||||||||||||||<br>Sbjct 58479916 AGGTCAACCAGAAACTTTACATGTGTGTGTGAGGGCCTCTCCTGCGGGAACGAGGACC 58479857<br><br>Query 295 -------AGGCCAGCAGTGTTTTTCTTCTCTGAGCATCAACGATGGCTTCCACGTCTACC 347<br>|||||||||||||||||||||||||||||||||||||||||||||||||||||</td></tr>
</table>

<table>
<tr><td></td><td>Sbjct 58479856 ACTGTGAAGGCCAGCAGTGTTTTTCTTCTCTGAGCATCAACGATGGCTTCCACGTCTACC 58479797<br>sgRNA ACACTGCTGGCCTTCACAGTGG Del:17<br><br>Query 348 AGAA----TGCTTTCTGGTTTATGAGCAGGGGAAGATGACGTGTAAGAccccccccGTCAC 403<br>||||    ||||||| |||||||||||||||||||||||||||||||||||| |||||||<br>Sbjct 58479796 AGAAGGGCTGCTTTCAGGTTTATGAGCAGGGGAAGATGACGTGTAAGACCCCGCCGTCAC 58479737<br>sgRNA AACCTGAAAGCAGCCCTTCTGG</td></tr>
<tr><td>4T1-KO-DAZL-1#</td><td>Query 309 TCATGCCAAACACCGTTTTTGT----AGAATAGATGTATGTCAATCGAATTTATATTTAA 364<br>||||||||||||||||||||||    |||| |||||   |   || | ||| |||||| |<br>Sbjct 50289132 TCATGCCAAACACCGTTTTTGTTGGAGGAATTGATGTTAGGGTATTGTATTCATATTTCA 50289073<br>sgRNA: CAAACACCGTTTTTGTTGGAGG Del: 4bp</td></tr>
</table>

| | |
|---|---|
| 4T1-KO-DAZL-2# | Query 310 ATGCCAAACACCGTTTTTG-TGGAGGAAATGATGTTAGGGTATTGTATTCATATTTCATT 368<br>||||||||||||||||||| |||||||| ||||||||||||||||||||||||||||||<br>Sbjct 50289130 ATGCCAAACACCGTTTTTGTTGGAGGAATTGATGTTAGGGTATTGTATTCATATTTCATT 50289071<br>sgRNA: CAAACACCGTTTTTGTTGGAGG Del: 1bp |
